# Delta-rate rhythm and voice familiarity modulate neural tracking in the infant brain

**DOI:** 10.64898/2026.09.10.750698

**Authors:** Isabelle Rambosson, Damien Benis, Francisca Barcos-Munoz, Claire Kabdebon, Leonardo Ceravolo, Didier Grandjean, Manuela Filippa

## Abstract

Delta-range rhythms (∼0.5–3.5 Hz) are a phylogenetically conserved tempo of biological communication, yet how the infant brain tracks them remains poorly understood. Thirty 6-month-olds heard syllable sequences produced by their mother or a stranger at a delta (2 Hz) or theta (4 Hz) rate, or at irregular intervals, while high-density EEG was recorded. Rhythmic sequences, at both rates, increased low-frequency power over frontotemporal sites. Over frontal sites, only the delta band showed rate-matched tracking: delta-band power was larger for delta-rate than for theta-rate or irregular sequences. This delta-rate-matched response strengthened across blocks. Voice familiarity modulated theta-band power: its increase for delta-rate over irregular sequences was larger for the stranger’s voice than the mother’s, whereas delta-band tracking was largely insensitive to voice identity. Cortical tracking at 6 months is tuned to the delta rate and shaped by voice familiarity, compatible with an early role of slow rhythms in temporal prediction.

## Introduction

Across the animal kingdom, many evolutionarily distant species communicate isochronously within a narrow tempo range of ∼0.5–3.5 Hz (∼2.7 Hz), suggesting a universal constraint rooted in the resonant properties of neural systems (*1, 2*). This temporal range closely corresponds to the delta frequency band and extends into human communication, where intonation and related dynamic shapes in speech establish a consistent low-frequency rhythm across languages (*3*). Converging phylogenetic evidence therefore identifies delta-band timing as a fundamental timescale of biological communication, paralleled by the predominance of delta activity during early human brain development.

In neonatal and infant EEG studies, delta activity is markedly elevated relative to later developmental stages and gradually decreases with cortical maturation and everyday experiences, indexing large-scale synaptic and thalamocortical reorganization (*4–6*). In neonates, delta oscillations often appear as delta brushes, hallmark patterns of early cortical activity associated with thalamocortical development and brain maturity (*4, 7*). This dominance persists, particularly during quiet sleep, in which slow-wave activity reflects sleep-dependent plasticity and homeostatic regulation (*8*). Beyond sleep, delta rhythms support early sensory and cognitive processing: from the first year of life, infant cortical activity robustly tracks the slow temporal structure of auditory input, including speech, facilitating attentional grouping and temporal parsing (*5, 9, 10*). Delta oscillations thus emerge as a foundational temporal scaffold for early brain function, such as social communication.

At the neural level, tracking rhythmic input is thought to rely on aligning endogenous neural oscillations to the temporal structure of external stimulation. Neural entrainment, in its strict sense, constitutes a key mechanism underlying this process: by aligning the phase of endogenous oscillations to the temporal structure of an external input, the brain creates cyclical fluctuations in excitability that selectively gate incoming information (*11–13*). This alignment is thought to operate in a frequency-specific manner: during auditory and speech processing, delta-band activity (∼0.5–3.5 Hz) tracks slower, prosodic-level temporal structure, whereas theta-band activity (∼4–8 Hz) tracks faster syllabic-level regularities (*11, 14*). Given the intrinsic dominance of delta rhythms in early development, neural tracking at this timescale may play a dominant role in infancy.

Speech constitutes a primary source of structured auditory input, and its processing relies on cortical tracking of temporal modulations predominantly in the delta and theta frequency bands (*9*), which closely mirror the temporal organization of natural speech. Within multi-time-resolution frameworks of speech processing, low-frequency oscillations, particularly delta, encode slow prosodic and phrasal structure and form the top level of a hierarchical system that organizes the faster theta and higher-frequency processing streams (*11, 15*). In infancy, cortical tracking is strongest in the delta band, suggesting that slow temporal structure provides a scaffold for early speech parsing (*16–18*). However, when 4- and 8-month-old infants listened to synthesized syllable strings with nested rhythmic units, cortical tracking was most robust at the syllable rate (5 Hz) and inconsistent at the slower rate of bisyllabic feet (2.5 Hz), across English-, French-, and Japanese-learning infants (*19*). Critically, such a system is already at play during pregnancy, allowing the fetus to process slow fluctuations in speech, especially in the mother’s voice (*20*), even though fine-grained segmental information remains largely inaccessible due to the filtering properties of the intrauterine environment (*21, 22*).

Importantly, the acoustic characteristics of infant-directed speech (IDS), including slower tempo, exaggerated prosody, and increased rhythmic regularity, enhance delta-band modulations relative to adult-directed speech (*15*). These features closely match infants’ endogenous oscillatory dynamics and promote stronger phase alignment in the delta-theta range (*15, 23, 24*). As a result, IDS appears to be optimally tuned to engage infants’ neural tracking mechanisms, thereby supporting speech segmentation, attention, and early language learning (*15, 23*). However, whether neural tracking at 6 months of age depends on the rate of the rhythmic input and on the familiarity of the voice remains to be established.

Auditory input in infancy is defined not only by its temporal structure but is also shaped by social experience, particularly through exposure to familiar voices. From birth, infants can discriminate their mother’s voice from that of a stranger (*25*). Indeed, neuroimaging studies reveal distinct patterns of cortical activation depending on speaker familiarity. The mother’s voice preferentially engages the left temporal regions associated with language processing, whereas unfamiliar voices more strongly recruit the right temporal regions involved in voice-specific processing (*26*). In full-term newborns, an unfamiliar voice also elicits early mismatch responses over frontocentral electrodes, consistent with detecting vocal novelty (*25*). Oscillatory activity further indicates differential engagement across frequency bands, with maternal voices eliciting broader spectral activation than those of strangers (*27*).

Speaker familiarity may thus modulate not only regional cortical activation but also the neural tracking of speech rhythm and prosody. The prosodic stress profile elicits stronger cortical responses in 7-month-old infants and more effectively drives neural oscillatory activity, supporting effective tracking of speech input (*15, 23*). This neural responsiveness to acoustic regularities, in turn, supports infants’ early speech recognition and word learning during the initial stages of language acquisition (*15, 23*). Among the rhythmic auditory inputs to which infants are exposed, the mother’s voice holds a uniquely privileged position and may therefore specifically modulate neural tracking, providing a direct rationale for including speaker familiarity as a key variable in the present study. Given the involvement of temporal and frontal regions in early voice processing (*25–27*), the present study focuses on the left frontal (LF), right frontal (RF), left temporal (LT), and right temporal (RT) regions as the main regions of interest.

The present study addresses these questions by examining neural tracking in 6-month-old infants using high-density electroencephalography (EEG; see Fig.1). Infants are exposed to rhythmic 4-syllable sequences produced by either their mother or a stranger at a slower, delta rate (2 Hz, one syllable every 500 ms) or a faster, theta rate (4 Hz, one syllable every 250 ms), as well as to a non-rhythmic control condition. This design allows us to test whether (a) rhythmicity modulates neural tracking relative to non-rhythmic sequences, (b) tracking is frequency-specific, and (c) speaker familiarity modulates tracking dynamics. In an additional exploratory analysis, conducted after inspection of the hypothesis-driven results, we examined whether neural tracking changed across successive blocks of exposure.

By characterizing these effects, the study aims to clarify how, at 6 months of age, the extraction of temporal regularities in auditory input depends on their rate, on repeated exposure, and on the familiarity of the voice producing them, as a potential step toward understanding the early foundations of temporal prediction.

## Results

To test our hypotheses, we first investigated whether rhythmic stimulation, presented at a delta (DEL) or a theta (THE) rate, elicits a distinct neural response compared to a non-rhythmic (NR) stimulation, and then its frequency-specificity, defined as the largest power in the corresponding frequency band (DEL > THE and DEL > NR for the delta frequency band; mirror pattern for theta). We further explored whether this pattern evolved with repeated exposure and examined its modulation by voice familiarity. Spectral power was analyzed across bilateral frontal and temporal, central, bilateral parietal, and posterior regions. All analyses included the eight regions of interest (ROIs) as a fixed factor in a single model per hypothesis (see Methods). The main text focuses on the four frontotemporal clusters (see Fig.1) documented above as implicated in early voice processing (*25–27*): left frontal (LF), right frontal (RF), left temporal (LT), and right temporal (RT). Complete omnibus results, post hoc results for the central, parietal, and posterior clusters, and effect sizes are provided in the Supplementary Materials (Tables S4–S14).

**Fig. 1.**
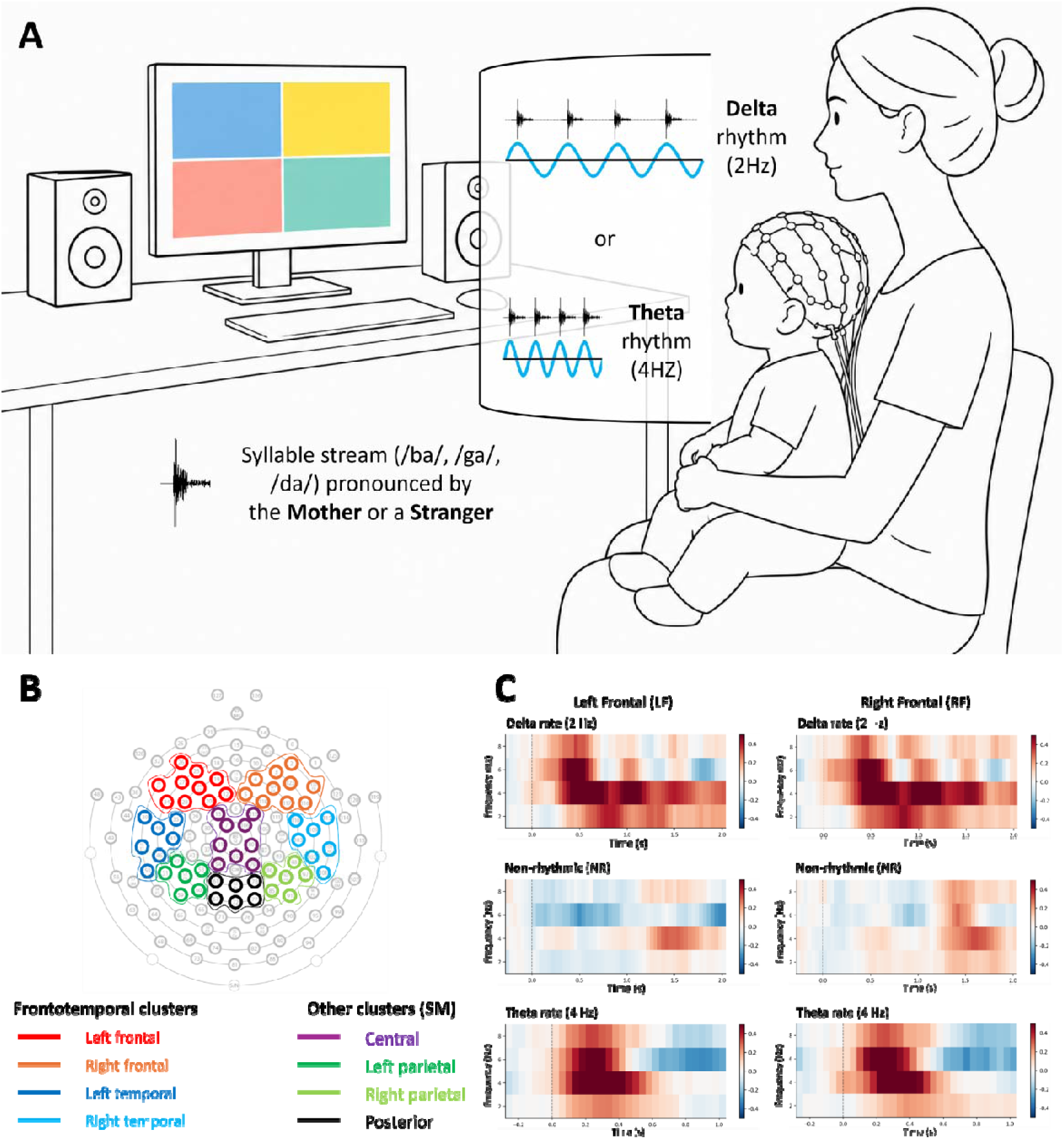
Experimental setup, regions of interest, and time–frequency responses. (B) Infants, seated on their mother’s lap, heard syllable sequences (/ba/, /ga/ or /da/) produced by their mother or by a stranger, presented at a delta rate (2 Hz) or a theta rate (4 Hz), or non-rhythmically (non-rhythmic or NR condition, not depicted). (B) Time–frequency power (2–8 Hz) over the left (LF) and right (RF) frontal clusters for delta-rate, non-rhythmic, and theta-rate sequences, averaged across the two voices. Values are power changes (dB) relative to a pre-sequence baseline (−250 to −50 ms), averaged across all artifact-free sequences, with no omissions from the 30 infants. Time 0 marks the start of the sequence window (2 s for delta-rate and non-rhythmic sequences, 1 s for theta-rate sequences).

### 1. Rhythmicity drives neural tracking in the infant brain

We first asked whether temporal regularity itself drives the infant brain’s response by comparing delta-rate (DEL) and theta-rate (THE) stimulations with non-rhythmic (NR) stimulation. Rhythmic stimulation (RHY) modulated low-frequency power across scalp regions (RHY: χ²(2) = 1054.29, *p* < .0001; RHY × ROI: χ²(14) = 113.60, *p* < .0001; Table S4).

Both rates contributed to this effect: in all four frontotemporal clusters, both delta-rate (DEL) and theta-rate (THE) stimulations elicited greater power than non-rhythmic (NR) stimulation (DEL vs. NR: est. = 0.218–0.268; THE vs. NR: est. = 0.104–0.197, all *ps* < .0001; Table S5).

### 2. Frequency-specific tracking emerges in the delta band only and builds up across blocks

We tested whether the effect of rhythmic stimulation differed between frequency bands (FRQ). Frequency-specific tracking requires each rate to drive the largest power in its own band: delta-rate sequences should dominate the delta band, and theta-rate sequences the theta band, each exceeding both the other rate and the non-rhythmic condition.

The effect of rhythmic stimulation differed between frequency bands (RHY × FRQ: χ²(2) = 39.28, *p* < .0001), and this interaction varied across regions (RHY × FRQ × ROI: χ²(14) = 53.46, *p* < .0001; Table S4).

The delta band met this criterion in the frontal clusters (see Fig.2). Delta-rate stimulation elicited greater delta-band power than non-rhythmic stimulation in all four clusters (est. = 0.119–0.246, *p*s ≤ .0002), and greater power than theta-rate stimulation in the left and right frontal clusters (est. = 0.080 and 0.073, *p*s ≤ .0201), but not in the temporal clusters (Table S6 and Fig.2).

The theta band did not meet this criterion. Theta-rate stimulation elicited greater theta-band power than non-rhythmic stimulation in all four clusters (est. = 0.149–0.224, *p*s < .0001), but delta-rate sequences elicited even more in the left and right frontal and left temporal clusters (THE vs. DEL: est. = −0.093 to −0.167, *p*s ≤ .0039), while the two rates did not differ in the right temporal cluster. Delta-rate sequences thus drove the largest response in the theta band, and no cluster showed theta-specific tracking.

In an exploratory analysis of the four successive blocks of each rhythmic condition (not planned a priori; see Materials and Methods), we examined whether the effects evolved with repeated exposure. Changes across blocks differed by rhythmic stimulation and frequency bands, and varied across regions (RHY × FRQ × BLK_ORD: χ²(2) = 27.76, *p* < .0001; RHY × FRQ × BLK_ORD × ROI: χ²(14) = 91.66, *p* < .0001; cf. Table S4).

Delta-band power increased across blocks for delta-rate sequences in all four frontotemporal clusters (slopes = 0.062–0.111, *ps* ≤ .0099; see Fig.2 and Table S7), but this increase exceeded that for non-rhythmic stimulation only in the right frontal cluster (DEL vs. NR = 0.077, *p* = .0047). Theta-band changes across blocks were heterogeneous across clusters.

**Fig. 2.**
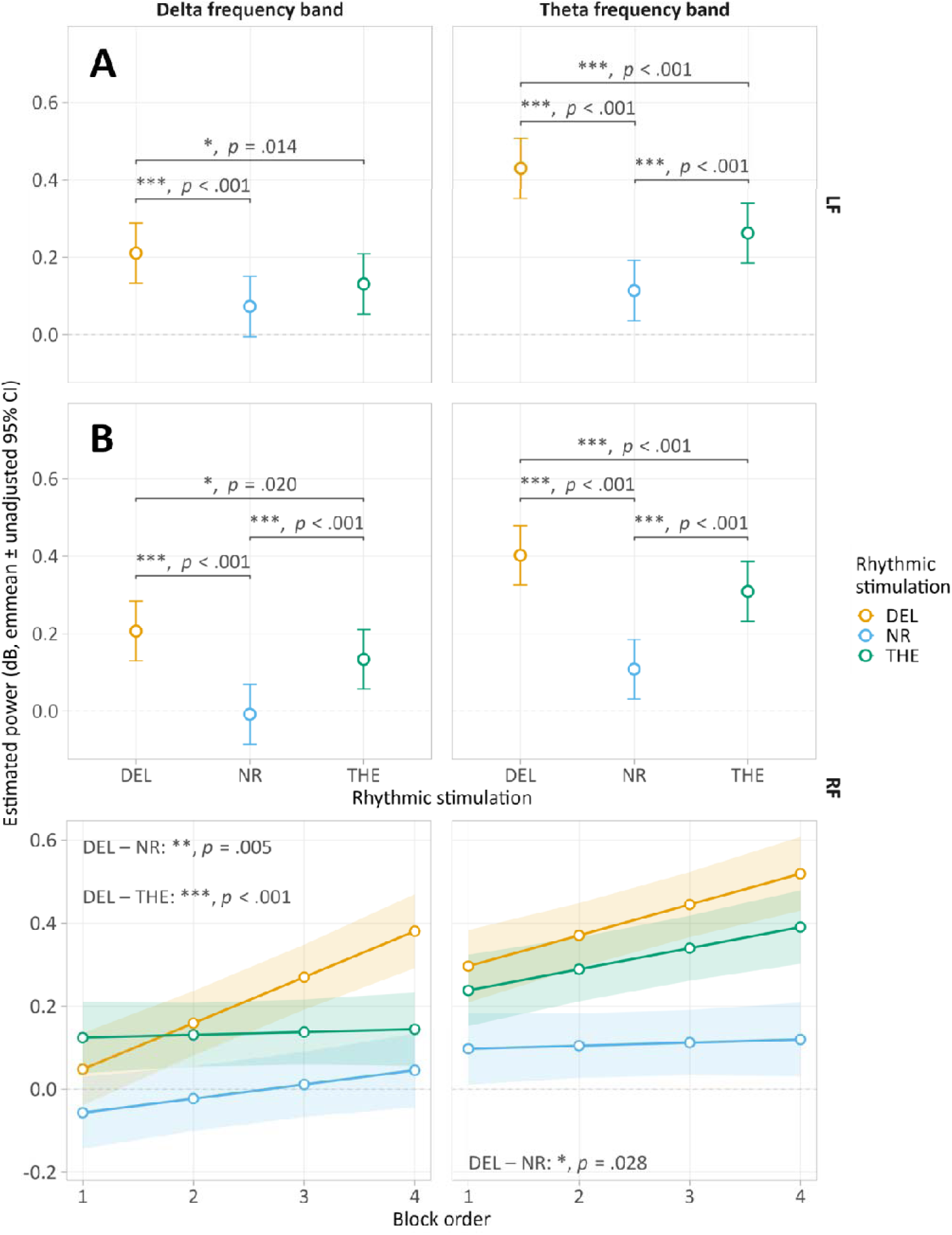
Delta-specific tracking over frontal clusters and its evolution across blocks. (**A**) Left frontal cluster (LF). Estimated marginal means (± unadjusted 95% CI) of baseline-corrected EEG power (dB) in the delta and theta frequency bands. Each panel displays the three rhythmic stimulations: delta rate (DEL, 2 Hz), non-rhythmic (NR, control), and theta rate (THE, 4 Hz). (**B**) Right frontal cluster (RF). Top: same representation as in (A). Bottom (exploratory analysis): model-estimated power (± unadjusted 95% CI) across the four successive blocks of each stimulation; lines show the linear effect of block order. Brackets (top rows) and labels (bottom row) indicate significant contrasts between means or between slopes, respectively, with p-values adjusted using the multivariate t method across the eight clusters (* *p* < .05, ** *p* < .01, *** *p* < .001). *N* = 30 infants.

### 3. Voice familiarity modulates theta-band responses

To test whether voice familiarity further shaped these responses, we compared sequences produced by the infant’s mother and by a stranger. Power differed between the two voices (VOICE: χ²(1) = 173.89, *p* < .0001), and the effect of rhythm across frequency bands depended on the voice (RHY × FRQ × VOICE: χ²(2) = 85.99, *p* < .0001), with differences across regions (RHY × FRQ × VOICE × ROI: χ²(14) = 36.54, *p* = .0009; Table S4).

The clearest voice effect was observed in the theta band, where the advantage of delta-rate over non-rhythmic stimulations was larger for the stranger’s voice than for the mother’s over the left and right frontal and left temporal clusters (Mother − Stranger: est. = −0.235 to −0.291, *ps* ≤ .0002; Table S8, panel 1). Over these clusters, the stranger’s voice also elicited a graded theta-band response (DEL > THE > NR; *ps* < .0001), which the mother’s voice did not show (Table S8, panel 3).

Voice familiarity had little influence in the delta band, except over the right temporal cluster, where the difference between delta- and theta-rate sequences was larger for the mother’s voice (est. = 0.170, *p* = .0286; Table S8, panels 1 and 2).

**Fig. 3.**
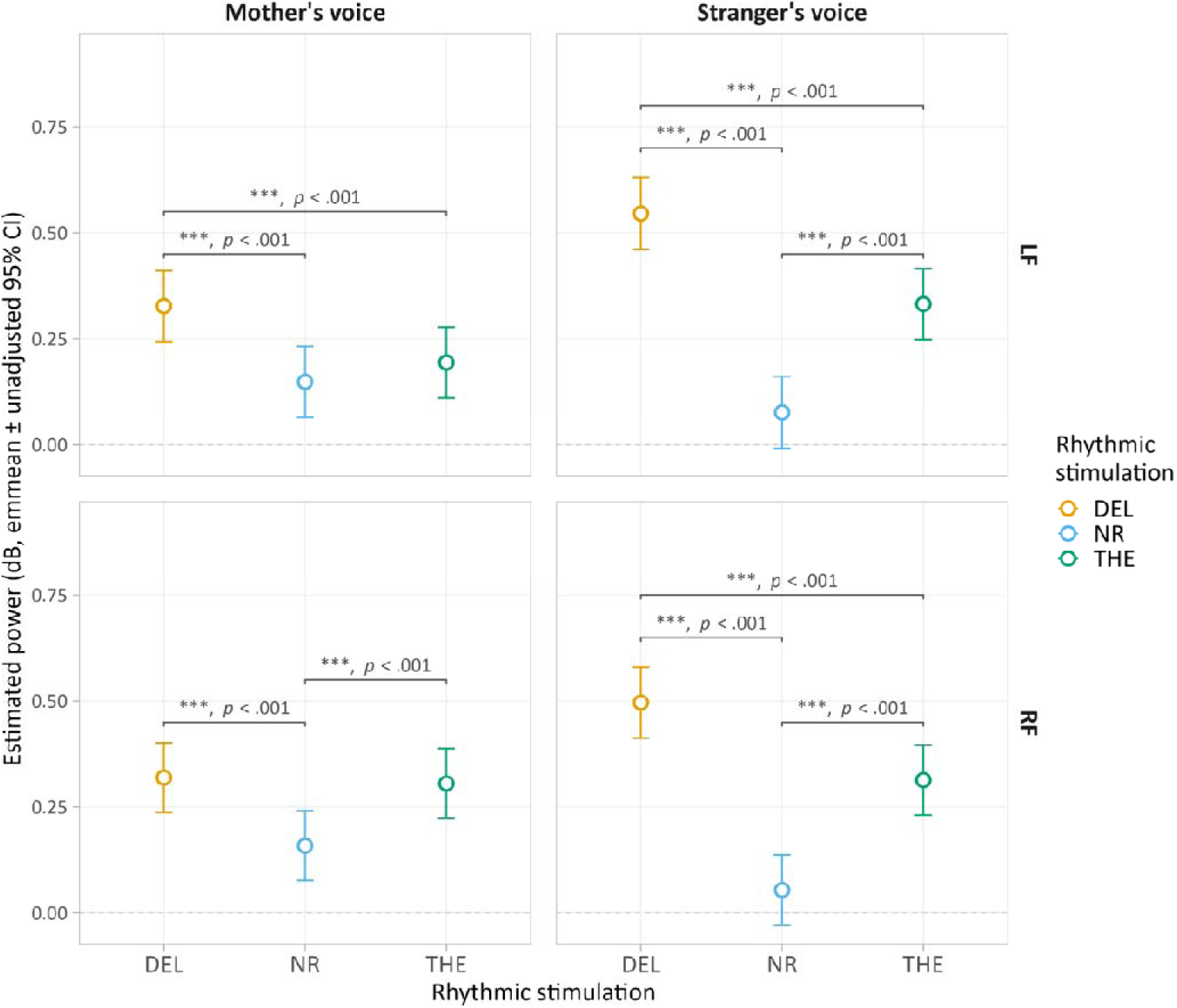
Frontal theta-band responses to rhythmic structure for the mother’s and stranger’s voices. Left frontal (LF, top) and right frontal (RF, bottom) estimated marginal means (± unadjusted 95% CI) of baseline-corrected theta-band EEG power (dB), shown separately for the mother’s voice (left) and the stranger’s voice (right). Each panel displays the three rhythmic stimulations: delta rate (DEL, 2 Hz), non-rhythmic (NR, control), and theta rate (THE, 4 Hz). Brackets indicate significant contrasts between means within each voice, with p-values adjusted using the multivariate t method across the eight clusters (*** *p* < .001). *N* = 30 infants.

## Discussion

Using EEG in 6-month-old term-born infants, we addressed three questions: whether rhythmicity drives neural tracking, whether this tracking is frequency-specific, and whether voice familiarity (i.e., mother’s vs. stranger’s voice) modulates it. An exploratory analysis further examined whether tracking evolved across successive blocks of exposure.

### 1. Rhythmicity drives broad frontotemporal neural tracking

The first key finding is that neural tracking is a functional mechanism at 6 months of age, as rhythmic auditory stimulation modulates oscillatory neural power relative to a non-rhythmic control condition across a broad frontotemporal network. This finding extends previous developmental EEG evidence showing that infants respond to rhythmic auditory input during the first year of life (*9, 10, 17*). Using a direct comparison with a non-rhythmic condition, we demonstrate that these responses are specifically driven by the temporal regularity of the auditory signal rather than by auditory stimulation per se. In isolating the contribution of temporal regularity, our findings support the hypothesis that neural tracking emerges early in auditory system development (*11, 12*) and suggest that temporal regularity may constitute an important organizing principle for auditory processing in infancy. These findings are consistent with the view that the infant brain actively extracts temporal regularities from its environment (*28*), a building block of more complex predictive abilities. That the immature brain already extracts rhythmic regularities and organizes them within a cortical hierarchy has been elegantly demonstrated in preterm neonates using non-linguistic auditory rhythms and beat/meter structures (*29,* see also *30, 31*). Building on this foundation, the present study shifts the question from *whether* the young brain detects rhythmic regularities to *whether and how* early experience shapes the way it does so, a shift we develop below through the frequency-specificity and voice-familiarity findings. Although we demonstrate this here through auditory stimulation, nothing in this mechanism inherently restricts it to the auditory domain.

### 2. Frequency-specific tracking in the delta band and its build-up across blocks

A central assumption of neural entrainment theory is that oscillatory power increases in the frequency band corresponding to the driving rhythmic stimulus’s rate (*11, 12*), reflecting a functional alignment between external inputs and endogenous neural activity (*34*). In adults, this frequency-specific processing is a characteristic feature of cortical speech processing (*11, 35*). The present results reveal that it is already in operation in six-month-old infants, albeit in a regionally constrained manner.

Delta-band frequency-specific tuning was found in two regions of interest: the left and right frontal clusters. This finding is consistent with an expanding body of research indicating that delta-band oscillations (∼0.5–3.5 Hz) are involved in the processing of slow, prosodic-level temporal regularities in both infants and adults (*9, 15, 17, 36*). Furthermore, in adults, frontal regions, including the inferior frontal gyri, are part of a network that processes auditory input and statistically builds up functional categorizations, allowing internal models to be updated (*37, 38*). Although scalp EEG does not allow precise localization of these sources, this role is compatible with the frontal distribution of the delta-specific responses observed here. Interestingly, this ability to selectively track the delta range is likely strengthened by the infant’s early exposure to IDS (characterized by a more prominent delta-band amplitude modulation profile than ADS, *9*). Such exposure may provide a sustained training signal for delta-band temporal processing from the first months of life (*15, 23, 24*) and constitutes one of the few stable, regular systems between pregnancy and birth. Notably, while Attaheri et al. (*9*) demonstrated preferential delta-band cortical tracking of sung speech, the present study experimentally manipulated the rate of syllable sequences. Furthermore, delta tracking has been shown to be present from birth (and probably during pregnancy) and to differentiate rhythmically distinct linguistic inputs (*17*), predicting later language outcomes, including vocabulary, phonology, and grammar, thereby highlighting the functional relevance of delta-band tracking for language acquisition even at play in early development (*9, 16, 39*). Its clinical relevance is further underscored by its sensitivity to developmental status: quantitative spontaneous delta power scales with gestational age and is disrupted in neonatal brain injury, making it a robust biomarker of early neural integrity and thalamocortical circuit development (*6, 40*). In this broader context, the selective delta-band tracking observed likely taps into a foundational temporal framework that, far from being a byproduct of immature neural activity, reflects a biologically grounded mechanism that organizes early brain function and potentially supports the emergence of speech perception and language development by updating the infant’s internal temporal models.

In an exploratory analysis, delta-band responses to delta-rate stimulation increased across blocks in all four frontotemporal clusters. However, this increase exceeded that for non-rhythmic stimulation, the better-matched control, only over the right frontal cluster, supporting a build-up specific to delta-rate regularity in this cluster alone. Sustained exposure to delta-rate syllable sequences thus appears to progressively enhance the corresponding low-frequency responses. This increase across blocks is compatible with several accounts, including familiarization, repetition enhancement, changes in attention or arousal, and the progressive refinement of temporal predictions, which the present design cannot disentangle.

By contrast, no theta-band frequency-specific tuning was observed in any electrode cluster. Although theta-rate stimulation increased theta-band power in all frontotemporal electrode clusters above the non-rhythmic control, this increase was exceeded by an even stronger delta-rate-driven theta response in the left and right frontal and left temporal clusters. This suggests that, at 6 months of age, delta-rate stimulation exerts a broadband influence that spills into the theta range rather than inducing strictly band-limited responses. This absence raises two non-exclusive interpretations. One interpretation is that the discrete, isolated syllables used in the present study do not accurately reflect the natural prosodic context in which syllabic-level theta tracking typically occurs (*11, 41*). A second interpretation relates to the rates used in the present study. In the present design, 2 Hz and 4 Hz both correspond to syllable rates, unlike studies in which the slower rates correspond to a higher-order prosodic level. Peter et al. (*19*) reported robust cortical tracking of syllable sequences at the syllable rate (5 Hz) in 4- and 8-month-old infants, with inconsistent tracking at 2.5 Hz, a rate that corresponds to bisyllabic feet. By contrast, when both rates were syllabic rates, as in the present study, the slower rate dominated responses in both frequency bands over the frontal clusters. Future studies using continuous, naturalistic IDS stimuli (see *9, 23 for examples*) will be better suited to disentangling the differential contributions of these alternatives.

### 3. Voice familiarity modulated theta-band responses

At 6 months of age, voice familiarity modulated theta-band power, most consistently in response to delta-rate stimulation. In the left and right frontal and left temporal clusters, delta-rate sequences boosted theta-band power more for the stranger’s voice than for the mother’s. Only the stranger’s voice also showed a stepwise increase in theta-band power from non-rhythmic to theta-rate to delta-rate sequences. This effect may reflect the greater novelty of the unfamiliar voice, which could engage attention or arousal more strongly, consistent with the role of theta activity in allocating attentional resources (*42, 43*). In the right temporal cluster, voice familiarity instead modulated delta-band power, with a larger difference between delta- and theta-rate sequences for the mother’s voice. These findings complement prior evidence from ERP and neural tracking paradigms showing differential responses to maternal and stranger voices in newborns and 7-month-old infants (*26, 44*).

Notably, the effects of voice familiarity were largely absent in the delta band response. Rather than reflecting a mere lack of sensitivity, this absence of a significant result may be theoretically informative. From birth onward, infants are extensively exposed to the delta-dominant modulation structure of infant-directed speech and can already track its speech envelope (*15, 24*). This exposure is expected to consolidate delta tracking into a robust cortical response driven primarily by the temporal envelope structure of speech rather than by voice-identity-specific features (*9, 11*). Within a predictive framework, a deeply embedded prior for delta-rate temporal structure could yield similarly low prediction errors for maternal and stranger voices (*45–47*), leaving little opportunity for differentiation based on familiarity. This aligns with hypotheses that exposure to IDS plays a fundamental role in developing the neural architecture for speech processing and temporal prediction (*15, 23*). In this context, the maternal voice, as the infant’s primary source of IDS, may have contributed to building this voice-general delta-band processing. Although infants preferentially respond to the maternal voice (*48, 49*) and learn more effectively from it (*50*), this preference does not appear to be expressed in the delta-band tracking.

### 4. Neural tracking and temporal prediction

Although the present study was designed to characterize neural tracking, this process represents a fundamental building block of predictive processing and can therefore be interpreted within this broader theoretical framework. In this framework, the brain is conceived as a predictive system that continuously generates and revises internal models to anticipate incoming sensory information, using prediction errors to update these models (*45–47, 51*). Because efficient processing of dynamic input requires anticipating not only what will occur but also when it will occur, temporal structure is central to this process, and endogenous rhythms aligned with biologically relevant timescales, such as the delta band, have been proposed as a substrate for organizing temporal predictions (*11, 13, 52*). Accounts differ, however, in how predictions and sensory signals interact. In the classical hierarchical view, top-down predictions carried by feedback connections are compared with bottom-up prediction errors (*53*). In this scheme, distinct oscillatory bands have been proposed to carry each stream, with delta-band activity conveying slower, prosodic-level temporal predictions and gamma-band activity signaling prediction errors (*14, 34*). A network-based account instead proposes that perception and its underlying internal model arise from the dynamic reorganization of distributed functional networks, in which predictions are carried by the temporal patterning of neural activity, such as oscillatory synchrony (*54, 55*). Predictive computations appear early in development (*56, 57*): 6-month-old infants show expectation-based feedback in the occipital cortex when an expected visual stimulus is omitted (*58*), and similar top-down responses have been reported in neonates (*59*).

Several aspects of the present findings are compatible with such an account. Rhythmic sequences, at both rates, elicited greater power than non-rhythmic sequences across frontotemporal clusters, and over the frontal clusters, delta-band power was selectively larger for delta-rate sequences. In the exploratory analysis, delta-band responses to delta-rate sequences increased progressively across blocks, but more steeply than responses to non-rhythmic sequences, only in the right frontal cluster, as expected if repeated exposure progressively refined a temporal model of the input. Similarly, the larger theta-band advantage of delta-rate over non-rhythmic sequences for the stranger’s voice could reflect the greater novelty, and thus the larger prediction error, associated with an unfamiliar speaker.

However, neural tracking can also arise without prediction, through the alignment of neural activity to the rhythm of the stimulus, and the present design does not allow these accounts to be distinguished. Paradigms that directly probe predictive mechanisms, such as omitting expected events within rhythmic sequences, will be needed to determine whether the neural tracking observed here reflects temporal prediction.

### 5. Limitations and recommendations for future studies

Several limitations of the present study should be acknowledged. First, the sample size (*n* = 30), although consistent with infant EEG studies in this field, limits statistical power, particularly for higher-order interactions. Interaction effects were small and should therefore be considered with caution until replicated in larger samples (*60*).

Second, using isolated syllables rather than continuous speech may not fully capture the natural temporal structure of IDS (*41, 61*). While this choice ensured strict experimental control over the stimulation rate, it may have limited the ecological validity of the tracking response and partly contributed to the lack of theta-band frequency specificity. Future studies using naturalistic IDS (e.g., nursery rhymes or spontaneous speech; see *9, 23*) would clarify whether this absence reflects a genuine developmental constraint or a stimulus-related limitation.

Third, although the present results demonstrate frequency-specific modulation of EEG spectral power, this pattern is consistent with frequency tagging and does not, by itself, constitute definitive evidence of genuine oscillatory entrainment. The operational definition of frequency specificity adopted in this study (cf. Supplementary Materials) was deliberately simplified relative to the strict definition of neural entrainment. True entrainment additionally requires evidence of post-stimulus oscillatory persistence and phase coherence with the stimulus envelope (*62, 63*), which the present power-based design cannot establish. Future studies should, therefore, incorporate post-stimulus silent periods and phase-based analyses (*64, 65*).

Fourth, because delta- and theta-rate stimulation differed in presentation rate, and because the second harmonic of the delta rate falls within the theta band, we cannot fully exclude that part of the observed effects reflects a generic difference between slower and faster stimulation, or a harmonic response, rather than band-specific tracking. Disentangling these accounts would require designs that include additional stimulation rates and should be addressed in future studies.

Fifth, the cross-sectional design of the present study precludes inferences about the developmental trajectory of these effects. Given the rapid, non-linear changes in oscillatory organization during the first year of life (*66, 67*), longitudinal designs tracking infants from 3 to 12 months would be essential to determine whether frequency-specific tuning and voice-familiarity effects strengthen or reorganize over this period.

Finally, the absence of a concurrent behavioral measure of voice preference (e.g., preferential looking or conditioned head turning) prevents a direct association between the observed neural patterns and behavioral sensitivity. Incorporating such measures in future studies would help determine whether these neural differences translate into differential attentional engagement or learning outcomes (see *48, 50 for examples*).

### 6. Conclusion

The present findings provide evidence that, by 6 months of age, the infant brain is already equipped with functional mechanisms for extracting and organizing temporal regularities in auditory input. Neural tracking was driven by rhythmicity across a broad frontotemporal network, with a clear primacy of delta-rate processing emerging in frontal regions. This preferential sensitivity to slow temporal regularities suggests that delta-band activity may provide a foundational temporal scaffold for early auditory processing, upon which the extraction of increasingly complex structures can develop. In any exploratory analysis, delta-band responses to delta-rate stimulation increased across exposure blocks, beyond the non-rhythmic control over the right frontal cluster, suggesting that this processing may remain sensitive to ongoing experience. Importantly, voice familiarity modulated theta-band responses, particularly to delta-rate stimuli, indicating that the processing of temporal structure is not independent of the social characteristics of the input. Together, these findings suggest that early neural tracking reflects an interplay between the extraction of temporal regularities, their experience-dependent refinement, and sensitivity to the characteristics of the auditory environment. As a fundamental building block of predictive processing, neural tracking may provide an early temporal scaffold upon which increasingly sophisticated predictive abilities develop. Although the present findings do not establish predictive mechanisms directly, they highlight the importance of investigating how early experience with rhythmic and socially relevant auditory input shapes the development of temporal expectations. More broadly, understanding how the infant brain extracts, prioritizes, and uses temporal regularities may provide important insights into the early foundations of speech perception and language development.

## Materials and Methods

### 1. Participants

Thirty-three 6-month-old term-born infants were recruited. Participants’ characteristics are presented in Supplementary Materials (Table S1). The exclusion criteria included the following: (i) neurological lesions (i.e., presence of a periventricular leukomalacia grade 2–4 or an intraventricular hemorrhage grade 3–4), (ii) suspicion of an early diagnosis of developmental disabilities (i.e., autism spectrum disorder), (iii) mothers with drug or alcohol abuse during pregnancy, or with a history of major depression and/or symptoms. The study was approved by the Cantonal Ethics Committee (BASEC No. 2022-02154) and was not preregistered. Written informed consent was obtained from parents in accordance with the Declaration of Helsinki.

Of these 33 participants, 3 were excluded due to fussiness or technical issues during the experimental session. Overall, data from 30 participants (mean age at test: 6.13 months; *SD*: 0.21; 12 females) were analyzed. Data from 3 participants were partial because the infants did not complete the entire experimental session.

### 2. Design, procedure, and stimuli

Approximately one week before the laboratory session, each mother recorded her voice at home with her own cell phone, following written instructions and an example recording. She produced the syllables /ba/, /ga/, and /da/, each repeated in three trains of four, as if addressing her infant in a calm moment. The infant was not present during the recording. The stranger’s voice presented to each infant was the voice of the mother of the previously tested infant, recorded with the same procedure. Individual syllables were then extracted from these recordings, resampled, faded in and out, and normalized (cf. Supplementary Materials for a full description of the stimulus processing procedure).

After the EEG was set up, infants and their mother or parents were invited to enter the experimental room. During the experiment, infants were seated on their parents’ laps, 1 m from a computer screen, in a Faraday cage. Auditory stimuli were presented at a sound pressure level of 65 dBA through two loudspeakers placed 1.5 m in front of the infant and at eye level. In contrast, visual stimuli were displayed on a 1920 × 1080 px computer screen. Both were delivered using PsychoPy (v2023.2.3; *68*); stimulus onset and offset triggers (10 ms pulses) were sent to the EEG amplifier via a parallel port to ensure precise synchronization with the EEG recording. Once the infant was quiet and awake, the experimental paradigm began with a 3-minute resting state, followed by the testing procedure. If the infant became restless, a short break was given; in some cases, additional time was allocated for feeding before continuing. However, if the infant could not return to a calm, awake state, the experimental session ended. To ensure consistency and optimal EEG signal throughout the session, the parent was asked not to move or interact with their infant during stimulus presentation. However, between experimental blocks, parents could soothe the infant if he/she became agitated; the use of a pacifier was permitted.

The testing procedure was organized into 12 blocks, each corresponding to one of the three rhythmic stimulus structures (delta, theta, or non-rhythmic control), with condition order pseudorandomized across blocks. In each block, a test phase of 8–10 trials followed 4– 6 familiarization trials. The paradigm was designed to address both neural tracking and temporal prediction: four test trials omitted a syllable, so that, depending on block length (12–16 trials), 25% to 33% of trials contained an omission. Omission trials, which address temporal prediction, are analyzed in a separate study [in prep]. The present study focuses on neural tracking during sequences that neither contained nor followed an omission (non-violation or NV trials).

Each trial consisted of four sequences of four syllables (16 syllables in total). Within a trial, all sequences repeated the same syllable (/ba/, /ga/ or /da/), produced by the same voice, using different recorded tokens whose order varied across sequences according to a predefined set of permutations. Syllable type and voice (the infant’s mother or a stranger) varied pseudorandomly across trials within each block. Sequences were presented at a delta rate (2 Hz, one syllable every 500 ms, sequence duration 1.66 s), at a theta rate (4 Hz, one syllable every 250 ms, sequence duration 0.90 s), or with non-rhythmic timing (NR, control). In NR sequences, the four syllable onsets were drawn at random within the same 2-s sequence slot as delta-rate sequences, with a minimum inter-onset interval of 250 ms and the last onset occurring no later than 1800 ms. NR sequences were generated anew for each infant under these constraints. Full timing parameters are reported in the Supplementary Materials.

Infants were also presented with geometric visual stimuli (of different colors), temporally matched to the onset of each auditory sequence. Audio-visual pairings were pseudorandomized across stimulus blocks and infants. This variability in syllabic and visual forms was intended to maintain the infant’s attention and to demonstrate that auditory rhythm is critical (rather than the specific characteristics of the sensory inputs).

#### a. Outcome measures

EEG data were recorded throughout the task using an Electrical Geodesics, Inc. (EGI)™ Geodesic Sensor Net (GSN) with a 128-channel cap. Most participants were recorded using an ANT Neuro eego™ amplifier; a minority were recorded using an EGI amplifier. The EGI net was used consistently across both systems to ensure data comparability. The experimental setup was optimized to ensure high-quality signal acquisition. Data were acquired at a sampling rate of 1000 Hz, and electrode impedances were maintained below 50 kω.

Before the experiment, parents’ socioeconomic status (SES; *69*) and maternal mental health were measured using the PSI (*70*) and the EPDS (*71*). Infants’ behavior was also assessed using the IBQ (*72*). Participants’ scores are presented in the Supplementary Materials (Table S1).

### 3. EEG data processing

Data processing was performed using MNE-Python (*73*), with six custom scripts executed sequentially.

Following import, electrode labels were mapped to the GSN-HydroCel-128 montage, behavioral logs were synchronized with EEG triggers, and events were defined at the onset of each four-syllable sequence. After removal of peripheral electrodes (105 channels retained), bad channels were interpolated, and the data were re-referenced to the common average and band-pass filtered (1.5–100 Hz, with a 50 Hz notch filter). Epochs were extracted relative to sequence onset, and ocular, cardiac, muscular, and sucking artifacts were removed using independent component analysis (ICA). Time-frequency representations (TFRs) were computed for each channel using multitaper convolution, converted to decibels, and expressed relative to a pre-stimulus baseline (−250 to −50 ms). Based on a priori hypotheses derived from the infant EEG literature, power was averaged within two canonical frequency bands (delta: 2–3 Hz and theta: 3–5 Hz) and within four consecutive post-stimulus windows (T1–T4) matching the syllable period of each condition (500 ms for delta-rate and non-rhythmic sequences, 250 ms for theta-rate sequences; cf. Supplementary Materials).

Artifacts were rejected in two stages based on peak-to-peak amplitude: a coarse threshold applied during epoching, followed by a data-driven threshold (median ± 5 *SD* of the amplitude distribution across participants, channels and epochs) applied channel by channel. On average, 7.9% of sequences were rejected, leaving 72–81 epochs per condition per infant (range = 23–97; cf. Supplementary Materials, Table S2).

To structure the analysis by scalp region, eight scalp ROIs were defined a priori based on existing literature: Left Frontal (LF), Right Frontal (RF), Left Temporal (LT), Right Temporal (RT), Central (CEN), Left Parietal (LP), Right Parietal (RP), and Posterior (POS), each comprising 6–10 electrodes (electrodes listed in the Supplementary Materials). Condition contrasts (rhythmic vs. non-rhythmic, mother vs. stranger) were examined using difference TFR maps computed for each ROI, centered on the frequency bands of interest (Delta: 2–3 Hz and Theta: 3–5 Hz).

For group-level analyses, TFR, power spectra, and behavioral data were concatenated across participants, organized by voice (mother, stranger) × rhythmic stimulation (DEL, THE, NR). Statistical analyses were conducted on single-epoch data, with each epoch × electrode × time window as an observation.

The complete pipeline and all parameter settings are described in the Supplementary Materials.

### 4. Statistical analysis

Models were fitted in R (v4.5.1; *74*) with the glmmTMB package (v1.1.13; *75*). The dependent variable (Power, in dB) was the spectral power of the EEG response to the syllable sequences, estimated from multitaper time-frequency representations, corrected relative to the pre-stimulus baseline, and averaged within the delta and theta bands, matching the stimulation rate of each condition. Only trials in which the rhythmic regularity of the sequence was preserved (i.e., NV trials; see Supplementary Materials) entered the analyses.

Because power was quantified at the stimulation frequency for each condition, increased power at these frequencies was interpreted as evidence of neural tracking of the auditory rhythm. For each hypothesis, a single model was fitted to the data of the eight scalp regions of interest (ROIs, cf. Supplementary Materials), with ROI as a fixed factor crossed with all other fixed factors, and with both frequency bands and the four post-stimulus windows (T1– T4) included as repeated observations, without a separate time factor. The random-effects structure was selected by AIC-based comparison of models including random intercepts for participant (PAT), electrode (CH), both, or neither. The model including random intercepts for both PAT and CH provided the best fit for all hypotheses (ΔAIC ≥ 50.1, cf. Supplementary Materials). Covariates were evaluated through a two-step procedure that combined AIC comparisons and likelihood ratio tests. Of the candidate covariates, only block order improved model fit (cf. Supplementary Materials). Block order (BLK_ORD) was included in all models a priori to control for session-wide habituation, and was operationalized as a continuous variable (1–12, reflecting each block’s overall sequential position across the session). In the exploratory block-order analysis, block order was realigned within each rhythmic condition (1–4), so that estimated marginal slopes reflect within-condition temporal dynamics rather than session-wide habituation. Three hypotheses were tested using Generalized Linear Mixed Models (GLMMs) with a Gaussian distribution and identity link function

1. the effect of rhythmic stimulation (RHY) on neural tracking.
2. the frequency-specificity (FRQ) of this effect.
3. the modulation of the rhythm-by-frequency interaction by voice familiarity (VOICE: mother vs. stranger).

Omnibus effects were assessed with Type II Wald χ² tests (car package, v3.1.3; *76*). As each term was tested once, in a single model including all ROIs, omnibus tests were not corrected for multiple comparisons. Post hoc contrasts were estimated using the emmeans package v2.0.0 (*77*). They were computed by ROI when the interaction between the term of interest and ROI was significant (RHY × ROI for hypothesis 1, RHY × FRQ × ROI for hypothesis 2, RHY × FRQ × VOICE × ROI for hypothesis 3), and averaged across ROIs otherwise.

Contrasts were corrected using the multivariate t adjustment (mvt, equivalent to Tukey’s HSD for complete pairwise comparisons; *78*), each family comprising all contrasts of a given set across the eight ROIs (24 contrasts), separately for each frequency band, and for each voice in the case of within-voice rhythm contrasts. Confidence intervals of contrasts were adjusted accordingly, whereas those of estimated marginal means were not. After the hypothesis-driven analyses, an exploratory extended model (RHY × FRQ × BLK_ORD × ROI, with block order realigned from 1 to 4) tested whether responses evolved across blocks. This model was not planned a priori but was added after inspection of the hypothesis-driven results and is therefore reported as exploratory. It was followed up only if hypothesis 2 had yielded post hoc contrasts, using estimated marginal slopes of block order (emtrends) with the same decision rule and correction.

Effect sizes were quantified at the model level using marginal and conditional R² (*79*), computed with the performance package v0.15.3 (*80*), and at the contrast level as unstandardized model-derived estimates with simultaneous 95% confidence intervals (cf. Supplementary Materials, Tables S9–S14).

Full model specifications, the model-selection procedure, covariate evaluation, residual diagnostics, and effect size computation are described in the Supplementary Materials.

## Data and materials availability

Raw EEG data are not publicly available. Participant consent, approved by the Cantonal Ethics Committee (BASEC No. 2022-02154), does not authorize transferring data to external repositories. De-identified data may be available to qualified researchers upon reasonable request to the corresponding author, subject to ethics committee approval. Analysis scripts (Python preprocessing pipeline and R statistical analyses) are available at https://doi.org/10.26037/yareta:d773de5ttffazptffb2tnjpgty.

## Author contributions

I.R., D.B., C.K., D.G. and M.F. conceived the study. I.R., D.B., L.C., D.G. and M.F. developed the methodology. I.R., D.B., F.B.-M. and M.F. carried out the investigation. I.R. performed the formal analysis and prepared the visualizations. L.C., D.G. and M.F. validated the results. D.B., L.C., D.G. and M.F. supervised the work. D.G. and M.F. acquired funding. I.R. and M.F. wrote the original draft. I.R., D.B., F.B.-M., L.C., D.G. and M.F. reviewed and edited the manuscript. D.G. and M.F. jointly supervised this work. All authors approved the final version of the manuscript.

## Acknowledgments

The authors thank the Arcade Sages-Femmes Association (midwives’ association, Geneva), and Dr. Marie Janaillac (pediatrician specialized in neonatology, Clinique des Grangettes) for assistance with participant recruitment, Emmanuelle Maillard (master’s student’s, University of Geneva), Léa Ghezzi (bachelor’s student, University of Geneva), Nina Luna Marbehant (bachelor’s student, University of Geneva) for assistance with data collection, and Maïté Fontela and Lilou Dehondt (master’s students, University of Geneva) for assistance with data collection, preprocessing, and processing of the data.

## References

1. G. Amichay, V. Balasubramanian, D. M. Abrams, A widespread animal communication tempo may resonate with the receiver’s brain. PLOS Biology 24, e3003735 (2026).

2. T. Piette, C. Cathcart, C. Barbieri, K. Martin Ming, D. Grandjean, B. Bickel, E. Déaux, A.-L. Giraud, Animal acoustic communication has a conserved optimal rhythm within the neural delta range. Plos Biology 24, e3003798 (2026).

3. M. Inbar, E. Grossman, A. N. Landau, A universal of speech timing: Intonation units form low-frequency rhythms. Proceedings of the National Academy of Sciences 122, e2425166122 (2025).

4. K. Whitehead, M. P. Laudiano-Dray, J. Meek, L. Fabrizi, Emergence of mature cortical activity in wakefulness and sleep in healthy preterm and full-term infants. *sleep* 41, zsy096 (2018).

5. E. Ünsal, R. Duygun, İ. Yemeniciler, E. Bingöl, Ö. Ceran, B. Güntekin, From infancy to childhood: A comprehensive review of event-and task-related brain oscillations. Brain Sciences 14, 837 (2024).

6. Y. Kitase, T. Hiroma, Y. Onishi, Y. Koyabu, S. Jones, A. Yoshino, S. Hayashi, H. Hayashi, R. Hayashi, S. Shiraki, C. Ishikawa, Y. Miyosawa, D. Zafer, A. Oba, F. Yamaide, K. Kashima, T. Shiohama, K. Fujii, T. Kubota, Spectral Power Analysis of Delta Waves in Neonatal Electroencephalography: A Tool for Assessing Brain Maturation and Injury. Cureus 17, e80680 (2025).

7. J. R. C. Conde, D. G. Barrios, C. G. Campo, N. L. G. González, B. R. Millán, A. J. Sosa, Visual and quantitative electroencephalographic analysis in healthy term neonates within the first six hours and the third day of life. Pediatric Neurology 77, 54–60. e51 (2017).

8. V. Schechtman, R. Harper, R. Harper, Distribution of slow-wave EEG activity across the night in developing infants. Sleep 17, 316–322 (1994).

9. A. Attaheri, A. Ni Choisdealbha, G. M. Di Liberto, S. Rocha, P. Brusini, N. Mead, H. Olawole-Scott, P. Boutris, S. Gibbon, I. Williams, C. Grey, S. Flanagan, U. Goswami, Delta- and theta-band cortical tracking and phase-amplitude coupling to sung speech by infants. Neuroimage 247, 118698 (2022).

10. A. Ni Choisdealbha, A. Attaheri, S. Rocha, P. Brusini, S. A. Flanagan, N. Mead, S. Gibbon, H. Olawole-Scott, I. Williams, C. Grey, P. Boutris, H. Ahmed, U. Goswami, Neural detection of changes in amplitude rise time in infancy. Dev Cogn Neurosci 54, 101075 (2022).

11. A.-L. Giraud, D. Poeppel, Cortical oscillations and speech processing: emerging computational principles and operations. Nature neuroscience 15, 511–517 (2012).

12. P. Lakatos, J. Gross, G. Thut, A new unifying account of the roles of neuronal entrainment. Current Biology 29, R890–R905 (2019).

13. P. Lakatos, G. Karmos, A. D. Mehta, I. Ulbert, C. E. Schroeder, Entrainment of neuronal oscillations as a mechanism of attentional selection. science 320, 110–113 (2008).

14. L. H. Arnal, D. Poeppel, A. L. Giraud, Temporal coding in the auditory cortex. Handb Clin Neurol 129, 85–98 (2015).

15. V. Leong, M. Kalashnikova, D. Burnham, U. Goswami, The temporal modulation structure of infant-directed speech. Open Mind 1, 78–90 (2017).

16. A. Ni Choisdealbha, A. Attaheri, S. Rocha, N. Mead, H. Olawole-Scott, P. Brusini, S. Gibbon, P. Boutris, C. Grey, D. Hines, I. Williams, S. A. Flanagan, U. Goswami, Neural phase angle from two months when tracking speech and non-speech rhythm linked to language performance from 12 to 24 months. Brain Lang 243, 105301 (2023).

17. M. C. Ortiz-Barajas, R. Guevara, J. Gervain, Neural oscillations and speech processing at birth. iScience 26, (2023).

18. M. Peña, E. Pittaluga, J. Mehler, Language acquisition in premature and full-term infants. Proceedings of the National Academy of Sciences 107, 3823–3828 (2010).

19. V. Peter, M. Kalashnikova, S. v. Ommen, R. Mazuka, D. Burnham, T. Nazzi, Cortical Tracking of Speech Across Rhythm Classes in English, French, and Japanese Learning 4 and 8 Month Old Infants. Developmental Science 29, e70271 (2026).

20. C. Nallet, J. Gervain, Neurodevelopmental preparedness for language in the neonatal brain. Annual Review of Developmental Psychology 3, 41–58 (2021).

21. D. Querleu, X. Renard, F. Versyp, L. Paris-Delrue, G. Crepin, Fetal hearing. Eur J Obstet Gynecol Reprod Biol 28, 191–212 (1988).

22. K. J. Gerhardt, R. M. Abrams, Fetal exposures to sound and vibroacoustic stimulation. J Perinatol 20, S21–30 (2000).

23. M. Kalashnikova, V. Peter, G. M. Di Liberto, E. C. Lalor, D. Burnham, Infant-directed speech facilitates seven-month-old infants’ cortical tracking of speech. Scientific reports 8, 13745 (2018).

24. V. Leong, U. Goswami, Acoustic-emergent phonology in the amplitude envelope of child-directed speech. PloS one 10, e0144411 (2015).

25. A. Adam-Darque, M. P. Pittet, F. Grouiller, T. A. Rihs, R. H.-V. Leuchter, F. Lazeyras, C. M. Michel, P. S. Hüppi, Neural correlates of voice perception in newborns and the influence of preterm birth. Cerebral cortex 30, 5717–5730 (2020).

26. M. Beauchemin, B. Gonzalez-Frankenberger, J. Tremblay, P. Vannasing, E. Martinez-Montes, P. Belin, R. Beland, D. Francoeur, A. M. Carceller, F. Wallois, M. Lassonde, Mother and stranger: an electrophysiological study of voice processing in newborns. Cereb Cortex 21, 1705–1711 (2011).

27. M. Filippa, D. Benis, A. Adam-Darque, D. Grandjean, P. S. Hüppi, Preterm infants show an atypical processing of the mother’s voice. Brain and Cognition 173, 106104 (2023).

28. J. Piaget, M. T. Cook, The origins of intelligence in children. (1952).

29. M. Edalati, M. Mahmoudzadeh, G. Ghostine, G. Kongolo, J. Safaie, F. Wallois, S. Moghimi, Preterm neonates distinguish rhythm violation through a hierarchy of cortical processing. Developmental Cognitive Neuroscience 58, 101168 (2022).

30. I. Rambosson, D. Benis, C. Kabdebon, D. Grandjean, M. Filippa, The origins of time: a systematic review of the neural signatures of temporal prediction in infancy. Developmental Cognitive Neuroscience, 101655 (2025).

31. A. Basirat, S. Dehaene, G. Dehaene-Lambertz, A hierarchy of cortical responses to sequence violations in three-month-old infants. Cognition 132, 137–150 (2014).

32. M. Edalati, F. Wallois, J. Safaie, G. Ghostine, G. Kongolo, L. J. Trainor, S. Moghimi, Rhythm in the premature neonate brain: Very early processing of auditory beat and meter. Journal of Neuroscience 43, 2794–2802 (2023).

33. M. Edalati, F. Wallois, G. Ghostine, G. Kongolo, L. J. Trainor, S. Moghimi, Neural oscillations suggest periodicity encoding during auditory beat processing in the premature brain. Developmental Science 27, e13550 (2024).

34. L. H. Arnal, A. L. Giraud, Cortical oscillations and sensory predictions. Trends Cogn Sci 16, 390–398 (2012).

35. A. Hyafil, A.-L. Giraud, L. Fontolan, B. Gutkin, Neural cross-frequency coupling: connecting architectures, mechanisms, and functions. Trends in neurosciences 38, 725–740 (2015).

36. A. Attaheri, D. Panayiotou, A. Phillips, A. Ni Choisdealbha, G. M. Di Liberto, S. Rocha, P. Brusini, N. Mead, S. Flanagan, H. Olawole-Scott, U. Goswami, Cortical Tracking of Sung Speech in Adults vs Infants: A Developmental Analysis. Front Neurosci 16, 842447 (2022).

37. S. Frühholz, D. Grandjean, Processing of emotional vocalizations in bilateral inferior frontal cortex. Neuroscience & Biobehavioral Reviews 37, 2847–2855 (2013).

38. D. Grandjean, Brain networks of emotional prosody processing. Emotion Review 13, 34–43 (2021).

39. A. Attaheri, A. Ni Choisdealbha, S. Rocha, P. Brusini, G. M. Di Liberto, N. Mead, H. Olawole-Scott, P. Boutris, S. Gibbon, I. Williams, C. Grey, E. O. M. Alfaro, C. Brough, S. Flanagan, U. Goswami, Infant low-frequency EEG cortical power, cortical tracking and phase-amplitude coupling predicts language a year later. PLoS One 19, e0313274 (2024).

40. L. Cornelissen, S.-E. Kim, J. M. Lee, E. N. Brown, P. L. Purdon, C. B. Berde, Electroencephalographic markers of brain development during sevoflurane anaesthesia in children up to 3 years old. British journal of anaesthesia 120, 1274–1286 (2018).

41. J. E. Peelle, M. H. Davis, Neural oscillations carry speech rhythm through to comprehension. Frontiers in psychology 3, 320 (2012).

42. J. F. Cavanagh, M. J. Frank, Frontal theta as a mechanism for cognitive control. Trends in cognitive sciences 18, 414–421 (2014).

43. I. C. Fiebelkorn, S. Kastner, A rhythmic theory of attention. Trends in cognitive sciences 23, 87–101 (2019).

44. S. Jessen, M. Orf, J. Obleser, Neural Tracking of the Maternal Voice in the Infant Brain. Journal of Neuroscience 45, (2025).

45. K. Friston, A theory of cortical responses. Philosophical transactions of the Royal Society B: Biological sciences 360, 815–836 (2005).

46. K. Friston, The free-energy principle: a unified brain theory? Nature reviews neuroscience 11, 127–138 (2010).

47. A. Clark, Whatever next? Predictive brains, situated agents, and the future of cognitive science. Behavioral and brain sciences 36, 181–204 (2013).

48. A. J. DeCasper, W. P. Fifer, Of human bonding: Newborns prefer their mothers’ voices. Science 208, 1174–1176 (1980).

49. A. J. DeCasper, M. J. Spence, Prenatal maternal speech influences newborns’ perception of speech sounds. Infant behavior and Development 9, 133–150 (1986).

50. R. van Rooijen, E. Bekkers, C. Junge, Beneficial effects of the mother’s voice on infants’ novel word learning. Infancy 24, 838–856 (2019).

51. K. Friston, The history of the future of the Bayesian brain. NeuroImage 62, 1230–1233 (2012).

52. G. Stefanics, B. Hangya, I. Hernádi, I. Winkler, P. Lakatos, I. Ulbert, Phase entrainment of human delta oscillations can mediate the effects of expectation on reaction speed. Journal of Neuroscience 30, 13578–13585 (2010).

53. R. P. Rao, D. H. Ballard, Predictive coding in the visual cortex: a functional interpretation of some extra-classical receptive-field effects. Nature neuroscience 2, 79–87 (1999).

54. A. K. Engel, P. Fries, W. Singer, Dynamic predictions: oscillations and synchrony in top–down processing. Nature Reviews Neuroscience 2, 704–716 (2001).

55. W. Singer, Recurrent dynamics in the cerebral cortex: Integration of sensory evidence with stored knowledge. Proceedings of the National Academy of Sciences 118, e2101043118 (2021).

56. A. Berger, M. I. Posner, Beyond Infant’s Looking: The Neural Basis for Infant Prediction Errors. Perspectives on Psychological Science 18, 664–674 (2023).

57. M. Köster, M. Langeloh, C. Michel, S. Hoehl, Young infants process prediction errors at the theta rhythm. NeuroImage 236, 118074 (2021).

58. L. L. Emberson, J. E. Richards, R. N. Aslin, Top-down modulation in the infant brain: Learning-induced expectations rapidly affect the sensory cortex at 6 months. Proceedings of the National Academy of Sciences of the United States of America 112, 9585–9590 (2015).

59. N. G. Xiao, C. E. Robertson, L. L. Emberson, Evidence of Top Down Sensory Prediction in Neonates Within 2 Days of Birth. Developmental Science 29, e70114 (2026).

60. P. E. Clayson, K. A. Carbine, S. A. Baldwin, M. J. Larson, Methodological reporting behavior, sample sizes, and statistical power in studies of event related potentials: Barriers to reproducibility and replicability. Psychophysiology 56, e13437 (2019).

61. J. Gross, N. Hoogenboom, G. Thut, P. Schyns, S. Panzeri, P. Belin, S. Garrod, Speech rhythms and multiplexed oscillatory sensory coding in the human brain. PLoS biology 11, e1001752 (2013).

62. J. Obleser, C. Kayser, Neural entrainment and attentional selection in the listening brain. Trends in cognitive sciences 23, 913–926 (2019).

63. B. Zoefel, S. Ten Oever, A. T. Sack, The involvement of endogenous neural oscillations in the processing of rhythmic input: More than a regular repetition of evoked neural responses. Frontiers in neuroscience 12, 95 (2018).

64. S. Van Bree, E. Sohoglu, M. H. Davis, B. Zoefel, Sustained neural rhythms reveal endogenous oscillations supporting speech perception. PLoS biology 19, e3001142 (2021).

65. G. Novembre, G. D. Iannetti, Tagging the musical beat: Neural entrainment or event-related potentials? Proceedings of the National Academy of Sciences 115, E11002–E11003 (2018).

66. E. V. Orekhova, T. A. Stroganova, I. N. Posikera, Theta synchronization during sustained anticipatory attention in infants over the second half of the first year of life. International Journal of Psychophysiology 32, 151–172 (1999).

67. P. J. Marshall, Y. Bar-Haim, N. A. Fox, Development of the EEG from 5 months to 4 years of age. Clinical neurophysiology 113, 1199–1208 (2002).

68. J. Peirce, J. R. Gray, S. Simpson, M. MacAskill, R. Höchenberger, H. Sogo, E. Kastman, J. K. Lindeløv, PsychoPy2: Experiments in behavior made easy. Behavior research methods 51, 195–203 (2019).

69. R. Largo, D. Pfister, L. Molinari, S. Kundu, A. Lipp, G. Due, Significance of prenatal, perinatal and postnatal factors in the development of AGA preterm infants at five to seven years. Developmental Medicine & Child Neurology 31, 440–456 (1989).

70. R. R. Abidin, Parenting Stress Index: Professional Manual (Psychological Assessment Resources, Odessa, FL, ed. 3, 1995).

71. L. Murray, A. D. Carothers, The validation of the Edinburgh Post-natal Depression Scale on a community sample. The British Journal of Psychiatry 157, 288–290 (1990).

72. M. A. Gartstein, M. K. Rothbart, Studying infant temperament via the revised infant behavior questionnaire. Infant behavior and development 26, 64–86 (2003).

73. A. Gramfort, M. Luessi, E. Larson, D. A. Engemann, D. Strohmeier, C. Brodbeck, R. Goj, M. Jas, T. Brooks, L. Parkkonen, MEG and EEG data analysis with MNE-Python. Frontiers in Neuroinformatics 7, 267 (2013).

74. R Core Team. (R Foundation for Statistical Computing, 2025).

75. M. E. Brooks, K. Kristensen, K. J. van Benthem, A. Magnusson, C. W. Berg, A. Nielsen, H. J. Skaug, M. Maechler, B. M. Bolker, glmmTMB balances speed and flexibility among packages for zero-inflated generalized linear mixed modeling. The R Journal 9, 378–400 (2017).

76. J. Fox, S. Weisberg, An R Companion to Applied Regression (Sage, Thousand Oaks, CA, ed. 3, 2019).

77. R. V. Lenth, J. Piaskowski. (CRAN, 2025).

78. A. Genz, F. Bretz, Computation of Multivariate Normal and t Probabilities (Springer, Berlin, Heidelberg, 2009), vol. 195.

79. S. Nakagawa, H. Schielzeth, A general and simple method for obtaining R2 from generalized linear mixed effects models. Methods in ecology and evolution 4, 133–142 (2013).

80. D. Lüdecke, M. S. Ben-Shachar, I. Patil, P. Waggoner, D. Makowski, performance: An R package for assessment, comparison and testing of statistical models. Journal of Open Source Software 6, 3139 (2021).

