## Supplementary Materials for "Delta-rate rhythm and voice familiarity modulate neural tracking in the infant brain"

### 1. Methods

Full details of the participants' characteristics, voice-recording treatment, EEG data acquisition and preprocessing, the analysis pipeline, and statistical analysis are provided below.

#### 1.1. Participants

| Participants | *N* = 30 |
|---|---|
| Gestational age (GA) at birth (weeks), *M* ± *SD* | 39.53 ± 1.31 |
| Birth weight (g), *M* ± *SD*. Partial data (*n* = 29)[a] | 3427.93 ± 524.45 |
| Age at test (months), *M* ± *SD* | 6.13 ± 0.21 |
| Sex: female / male | 12 / 18 |
| EPDS (*1*) score, *M* ± *SD* | 6.47 ± 3.37 |
| IBQ (*2*) score, *M* ± *SD* | |
| - Activity level | 4.09 ± 0.79 |
| - Distress to limitations | 3.39 ± 0.66 |
| - Distress and latency to intense stimulus | 2.45 ± 0.82 |
| - Duration of orienting | 3.72 ± 1.09 |
| - Smiling and laughter | 4.85 ± 0.94 |
| - Soothability | 5.32 ± 1.05 |
| PSI (*3*) score, *M* ± *SD* | 66.43 ± 11.55 |
| SES (*4*) mother's score, *M* ± *SD* | 1.33 ± 0.88 |
| SES (*4*) father's score, *M* ± *SD*. Partial data (*n* = 29)[b] | 2.04 ± 1.28 |

***Table S1.*** **Characteristics of the participants.**
[a] The weight of one participant was missing.
[b] The SES score of one participant's father was missing.

#### 1.2. Voice Recording and Stimulus Processing Procedure

Mothers received written instructions and an example recording by e-mail, and returned their recording approximately one week before the laboratory visit. Recordings were made at home, in a quiet room, using the mother's own cell phone held at a constant distance of 10 cm from the mouth. Each mother produced the syllables /ba/, /ga/, and /da/, each in three consecutive trains of four repetitions (12 tokens per syllable), following the example recording and as if addressing her infant in a calm moment, to elicit an infant-directed register. Recordings were not supervised by the experimenters, and the infant was not present. The stranger's voice for each infant was the recording of the mother of the infant tested immediately before, so that all voices were recorded under identical conditions and each stranger's voice was unfamiliar to the infant. For the first infant tested, the stranger's voice was that of a mother who had taken part in the pilot study and was recorded using the same procedure.

Syllables (/ba/, /ga/, /da/) were extracted from each mother's voice recording using Audacity (v3.3–3.7). Each syllable was cut to a total duration of 150 ms, starting 20 ms before vowel onset to include the syllable attack. Audio files were exported in mono WAV format (48,000

Hz, signed 16-bit PCM). Recordings were then resampled to 44,100 Hz using the Python library librosa. Fade-in and fade-out ramps of 20 ms each were applied in Praat using the Vocal Toolkit. Peak amplitude normalization was then applied, followed by intensity scaling to 70 dB. For each syllable type, the 8 best consecutive syllables out of 12 were selected based on visual and auditory inspection in Audacity, retaining either syllables 1–8 or 5–12.

#### 1.3. Trial Structure and Stimulus Timing

Each trial comprised four 4-syllable sequences (16 syllables). Within rhythmic sequences, syllable onsets were separated by 500 ms (delta rate; 502 ± 10 ms) or 250 ms (theta rate; 251 ± 10 ms), yielding sequence durations of 1.66 s and 0.90 s, respectively. Non-rhythmic sequences were generated anew for each infant by drawing the four syllable onsets at random (1-ms resolution) within the same 2-s sequence slot as delta-rate sequences, with a minimum inter-onset interval of 250 ms and the last onset occurring no later than 1800 ms after slot onset. This yielded inter-onset intervals of approximately 240–1090 ms and sequence durations of 0.97–1.94 s, and all syllables ended within 1.95 s of slot onset. Consecutive sequences within a trial were separated by silent intervals of 1.39 s (delta rate), 1.14 s (theta rate), and 1.1–2.5 s (non-rhythmic), and successive trials by approximately 2 s.

#### 1.4. Data Importation, Event Reconstruction, and Session Splitting (script 1)

Raw EEG data were imported into MNE-Python using the BrainVision reader (mne.io.read_raw_brainvision). Electrode labels were renamed to the EGI E-notation (E1-E128), and the standard GSN-HydroCel-128 montage was applied. All remaining auxiliary bipolar channels were removed.

Behavioral data were imported from the task log file and synchronized with the EEG triggers via a trigger-matching algorithm that iteratively removed unmatched rows until the two sequences aligned. The resulting merged table encoded, for each trial: stimulus identity, voice (Mother vs. Stranger), rhythmic stimulation (DEL, THE, or NR), violation type (NV: non-violation, BV: before violation, DV: during violation, or AV: after violation), and the timing of each stimulus within the sequence. Position labels (1–4) identifying the ordinal position of each stimulus within its sequence were assigned by aligning trial-level event timestamps with enclosing sequence boundaries. A final alignment step matched the all-session behavioral array to the participant's ICA-epoch CSV using a composite key (sample timestamp and stimulus identity), producing a behavioral array strictly synchronized with the retained epochs.

Event structures were constructed at the sequence level, marking the onset of each four-syllable sequence in accordance with the behavioral log. Sequence-level validity was verified by checking inter-onset intervals against expected rhythms (Delta: 500 ms, Theta: 250 ms) and by requiring exactly 16 stimuli per trial (four 4-syllable sequences). The continuous recording was divided into session files by segmenting into blocks of five meta-sequences; each segment was saved as an independent .fif file with its event arrays and behavioral metadata.

#### 1.5. Preprocessing, Epoching, and ICA (script 2)

A standard set of peripheral electrodes was removed (E1, E8, E25, E32, E43, E48, E49, E56, E63, E68, E73, E81, E88, E94, E99, E107, E113, E119, E120, E125, E126, E127, E128), leaving a 105-channel montage. Noisy electrodes were identified by visual inspection of the raw data and the power spectral density (Welch method, 0–150 Hz) and were manually listed for each participant. Identified bad channels were interpolated using spherical spline interpolation, and data were re-referenced to a common average.

A bandpass filter (1.5–100 Hz, zero-phase FIR, Hamming window, firwin design) was applied, followed by a 50 Hz notch filter. Epochs were extracted at the sequence level (-2 to +7 s) around each sequence onset, and a peak-to-peak amplitude threshold of 5000 µV was applied during epoching to initially reject strong movement artifacts. Epochs were resampled to 250 Hz.

ICA was performed on the sequence-level epoch set using the extended Picard algorithm (n_components = 15, random_state = 20, max_iter = 10,000, decim = 4). Components were visually inspected using topographic maps and time-course plots; components that clearly reflected ocular (blinks, saccades), cardiac, or muscular artifacts were removed.

#### 1.6. Cross-Session Merging (script 3)

ICA-cleaned epoch files from multiple recording sessions of the same participant were concatenated using mne.concatenate_epochs. Behavioral metadata arrays were aligned with the retained epochs using each epoch's drop_log, ensuring one-to-one EEG-behavioral correspondence. Merged epoch files and behavioral arrays were saved as participant-level .fif and .npy files.

#### 1.7. Condition-Level Database Construction and Artifact Rejection (script 4)

For each participant, merged sequence-level epochs were band-pass filtered (1.5–30 Hz). TFRs were computed from unfiltered (broadband) epochs using multitaper convolution (frequencies 2–98 Hz in 2 Hz steps). For frequencies ≤ 40 Hz, n_cycles = f/2 and time_bandwidth = 2; for frequencies > 40 Hz, n_cycles = f/5 and time_bandwidth = 6. TFR values were log-transformed ($10 \times \log_{10}$).

Artifact rejection was carried out in two stages, based on peak-to-peak amplitude computed from the time-domain EEG signal. First, an initial peak-to-peak threshold of 5000 µV was applied during epoching. Second, a data-driven threshold was derived from the peak-to-peak amplitude distributions across all participants, channels, and epochs (-500 to +2000 ms window), after excluding non-physiological values (<1 µV or >1000 µV) from this distribution (the threshold was set at the median ± 5 *SD*; a deliberately liberal criterion chosen to maximize epoch retention given the inherently noisy nature of infant EEG data). This threshold was applied channel by channel to the time-domain signal, and any epoch exceeding it on a given channel was excluded from that channel's data, including the corresponding time-frequency representations, prior to averaging. Epochs were sorted by rhythmic stimulation (DEL, THE, NR), voice (Mother, Stranger, or combined), and violation type (NV/BV combined as "NV", DV, or AV). Single-trial TFR data were retained for statistical analysis, and per-channel averages across epochs were computed for each participant × condition combination for visualization only.

Because some infants did not complete all blocks, the number of sequences presented varied across infants (mean = 77–86 per condition, range = 37–103). On average, 7.9% of sequences were rejected as artifacts (6.4–10.2% across conditions), about 7% at the global amplitude-threshold stage and about 1% at the electrode level. This left 72–81 epochs per condition per infant (*SD* = 12–17, range = 23–97, Table S2), corresponding to 242–535 epochs per infant across conditions (median = 497). The slightly lower number of epochs in stranger-voice conditions reflected fewer presented sequences.

| Condition | Presented sequences M [min–max] | Retained epochs M (*SD*) [min–max] | Rejection rate % M (*SD*) |
|---|---|---|---|
| **DEL, Mother** | 85.9 [51–97] | 80.5 (14.2) [42.7–94.0] | 6.4 (8.1) |
| **DEL, Stranger** | 76.9 [50–94] | 71.7 (11.6) [49.7–89.5] | 6.9 (8.6) |
| **THE, Mother** | 86.1 [44–103] | 77.7 (16.7) [36.0–93.0] | 10.2 (10.5) |

| | | | |
|---|---|---|---|
| **THE, Stranger** | 83.6 [41–96] | 77.0 (16.7) [33.0–95.9] | 8.0 (8.5) |
| **NR, Mother** | 85.0 [39–97] | 78.0 (15.7) [27.6–96.8] | 8.8 (9.1) |
| **NR, Stranger** | 79.1 [37–95] | 72.4 (14.3) [23.4–90.8] | 8.9 (9.5) |

***Note.*** *DEL = delta rhythmic stimulation; THE = theta rhythmic stimulation; NR = non-rhythmic stimulation; voice familiarity (Mother, Stranger)*

***Table S2.*** **Number of presented sequences, retained epochs, and rejection rates per condition.**
Values are computed per infant, then summarized across the 30 infants. Retained epochs are averaged across electrodes, since artifact rejection was applied electrode by electrode. Presented sequences include non-violation sequences (NV and BV) from analyzed sessions only. DEL = delta rate, THE = theta rate, NR = non-rhythmic.

#### 1.8. Group-Level Database Merging (script 5)

Per-participant condition files were merged into group-level databases by iterating over all participants and appending single-trial data along the trial axis for each channel. Separate group databases were assembled for raw broadband, TFR, and behavioral data, organized by voice × rhythm condition, and saved as group-level .joblib files.

#### 1.9. Time-Frequency ROI Database Construction (script 6)

For each experimental condition, group-level TFR and raw epoch data were reloaded and subjected to a final round of artifact rejection using the same peak-to-peak threshold as in script 4 (median ± 5 *SD*). The epoch window was cropped to -1 to +2 s. After artifact removal, a pre-stimulus baseline correction was applied to single-trial TFR data: mean power in the -250 to -50 ms window was subtracted for each frequency bin across surviving epochs, capturing the pre-stimulus expectancy period.
Two canonical frequency bands were defined: Delta (2–3 Hz) and Theta (3–5 Hz). Five time windows relative to stimulus onset were defined: a pre-stimulus baseline (–250 to –50 ms) and four consecutive post-stimulus windows (T1–T4). For delta-rate sequences, the windows matched the syllable period (0–500, 500–1000, 1000–1500 and 1500–2000 ms after sequence onset), and for theta-rate sequences, the shorter syllable period (0–250, 250–500, 500–750 and 750–1000 ms). Because syllable onsets in non-rhythmic sequences were irregular, syllable-locked windows could not be defined, and non-rhythmic sequences were analyzed with the same four 500-ms windows as delta-rate sequences (0–2000 ms), which covered the entire sequence. For each surviving epoch, channel, frequency band, and time window, mean TFR power was stored as a row in a CSV file, along with condition labels (rhythmic stimulation, voice, frequency band, channel name, behavioral pattern, block order). Baseline-corrected TFR epochs were also averaged across trials and cropped to a final analysis window (-250 ms to +2 s for DEL and NR; -250 ms to +1 s for THE).
Eight scalp ROIs, each comprising 6–10 electrodes were defined a priori based on existing literature (*5-7*): Left Frontal (E12, E19, E20, E23, E24, E26, E27, E28, E33, E34), Right Frontal (E2, E3, E4, E5, E116, E117, E118, E122, E123, E124), Left Temporal (E35, E39, E40, E41, E45, E46, E50), Right Temporal (E101, E102, E103, E108, E109, E110, E115), Central (E7, E13, E31, E54, E55, E79, E80, E106, E112), Left Parietal (E47, E51, E52, E53, E59, E60), Right Parietal (E85, E86, E91, E92, E98, E97), and Posterior (E61, E62, E67, E78, E72, E77). Condition contrasts were visualized as difference TFR maps across all ROIs.

#### 1.10. Statistical Analysis

This section details the statistical approach summarized in the main text (Materials and Methods, Statistical analysis). Statistical analyses were performed in R (v4.5.1; *8*). Mixed models were fitted with glmmTMB (v1.1.13; *9*). Estimated marginal means, slopes, and contrasts were derived with emmeans (v2.0.0; *10*), and p-values and confidence intervals were adjusted for multiple comparisons with the multivariate t method (*11*). Type II Wald $\chi^2$ tests were computed with car (v3.1.3; *12*), marginal and conditional $R^2$ with performance (v0.15.3; *13*), and residuals were checked with DHARMa (v0.4.7; *14*). Figures were drawn with ggplot2 (v4.0.0; *15*). All models used the same outcome, Power (in dB), defined as the spectral power of the EEG response to the syllable sequences, derived from multitaper time-frequency decompositions and expressed relative to the pre-stimulus baseline. Analyses were restricted to sequences not preceded by a violation trial, i.e., NV trials (see section Trial Structure and Stimulus Timing). For each hypothesis, a single model was fitted to the data of the eight regions of interest (ROIs): left frontal (LF), right frontal (RF), left temporal (LT), right temporal (RT), central (CEN), left parietal (LP), right parietal (RP), and posterior (POS), each defined by a predetermined set of electrodes, with ROI included as a fixed factor crossed with all other fixed factors. Data were filtered to retain only the delta and theta frequency bands and the four post-stimulus windows (T1–T4), which entered the models as repeated observations, without a separate time factor.

**Model selection.** For each hypothesis, the random-effects structure was first determined through AIC-based model comparison. Four candidate models were evaluated, differing in whether participant identity (PAT) and electrode (CH) were included as random intercepts: a fixed-effects-only model, models with PAT alone or CH alone, and a model combining both random intercepts (Power ~ fixed effects + (1|PAT) + (1|CH)). Candidate models included the fixed effects of each hypothesis crossed with ROI. The model combining both random intercepts yielded the lowest AIC for all hypotheses (ΔAIC = 50.1–50.2 relative to PAT alone, and > 4800 relative to CH alone or no random effect; Table S3) and was retained.

| Model | No random effect ΔAIC | (1\|CH) ΔAIC | (1\|PAT) ΔAIC | (1\|PAT) + (1\|CH) ΔAIC |
|---|---|---|---|---|
| **1** | 4919.9 | 4870.3 | 50.1 | 0 (best) |
| **2** | 4920.3 | 4870.7 | 50.1 | 0 (best) |
| **3** | 4909.1 | 4859.5 | 50.1 | 0 (best) |
| **Extended model (exploratory)** | 4891.1 | 4841.4 | 50.2 | 0 (best) |

***Table S3.*** **Random-effects selection: AIC difference (ΔAIC) relative to the best model for each hypothesis.**

Following random-effects selection, covariates were evaluated using a two-step procedure that combined AIC comparisons and likelihood ratio tests (LRTs). An initial model containing only the fixed effects of interest (crossed with ROI) was compared against thirteen augmented models, each adding one covariate: block order (BLK_ORD), infant sex (SEX), age at testing (AGE), maternal postnatal depression score (EPDS), and six subscales from the Infant Behavior Questionnaire (IBQ: activity level [IBQ_AL], distress to limitations [IBQ_DL], distress and latency to intense stimulus [IBQ_DI], duration of orienting [IBQ_DO], smiling and laughter [IBQ_SL], soothability [IBQ_S]), the Parenting Stress Index total score (PSI), and parents' socioeconomic status (SES_M and SES_F). One missing paternal SES score was imputed with the sample median. A covariate was retained in the final model when it both minimized AIC and produced a significant LRT ($\chi^2$ test, $p < .05$). Across the three hypothesis-driven models, only block order (BLK_ORD) met both criteria, yielding the lowest AIC (ΔAIC = 4.97, 4.98 and 3.38 relative to the model without covariate for Models 1, 2 and 3, respectively) and a significant LRT ($\chi^2(1) = 6.97$, 6.98 and 5.38, $p$ = .008, .008 and .020, respectively). None of the other covariates improved model fit (all LRT $ps \geq .29$), and they were therefore not retained. This result was consistent with the a priori inclusion of BLK_ORD in all models to control for session-wide habituation. Block order was operationalized as a continuous variable reflecting overall session-wide block position and habituation (1–12), whereas in the exploratory slope analysis, it was realigned within each rhythmic structure (1–4), so that estimated marginal slopes capture within-condition temporal dynamics.

**Hypothesis-specific models.** Three hypotheses were tested, each addressed with a dedicated Generalized Linear Mixed Model (GLMM) with a Gaussian distribution and identity link, fitted using glmmTMB on the data of the eight ROIs (6.59 million observations). All models included ROI as a fixed factor crossed with the factors of interest, block order (BLK_ORD) as a covariate, and random intercepts for participant (PAT) and channel (CH). Model residuals were inspected using the DHARMa package (250 simulations) to verify the absence of systematic misfit (uniformity, dispersion, and outlier tests). Scaled residuals closely followed the expected uniform distribution, with no evidence of over- or underdispersion (dispersion ratio = 1.00, $p$ = .992 for all models, including the exploratory extended model). Because of the very large number of observations, Kolmogorov–Smirnov and outlier tests were significant, but deviations were negligible in magnitude (D = 0.016, 1.0% vs. 0.8% expected outliers for all models).

- *Model 1* examined the effect of rhythmic stimulation on neural tracking: Power ~ RHY × ROI + BLK_ORD + (1|PAT) + (1|CH), where RHY (rhythm condition: DEL, THE, NR) was the fixed effect of interest, crossed with ROI (eight levels).
- *Model 2* tested whether the effect of the rhythmic stimulation differed between frequency bands: Power ~ RHY × FRQ × ROI + BLK_ORD + (1|PAT) + (1|CH), where FRQ (Delta, Theta) was added and its interaction with RHY examined. Frequency-specific tracking was operationally defined as significantly greater delta power in the DEL condition compared to both the THE condition (frequency-specificity criterion: DEL > THE) and the NR condition (tracking criterion: DEL > NR). The mirror pattern was predicted for the theta band (THE > DEL and THE > NR). This simplified criterion does not require the absence of elevation in the non-target band (i.e., Delta: DEL > NR > THE and Theta: THE > NR > DEL. Neither this criterion nor the full ordering pattern establishes neural entrainment in its strict sense, which would additionally require evidence of oscillatory persistence after stimulus offset and of phase alignment with the stimulus (see Limitations). The simplified criterion reflects the known immaturity of frequency-specific oscillatory organization in early infancy, including broadband cortical responses, incomplete myelination, and less distinct spectral differentiation between the delta and theta ranges than in adults (*16-21*). This approach is consistent with prior infant EEG studies examining neural tracking of beat, meter, and speech (see *20, 22 for examples*) and avoids applying adult-derived spectral segregation standards to a population for which such standards have not yet been fully established. After the hypothesis-driven analyses, an exploratory extended model tested whether responses evolved across blocks, with block order included as an interaction term: Power ~ RHY × FRQ × BLK_ORD × ROI + (1|PAT) + (1|CH), block order being realigned from 1 to 4 within each rhythmic structure.
- *Model 3* investigated whether voice familiarity modulated the rhythm-by-frequency effect: Power ~ RHY × FRQ × VOICE × ROI + BLK_ORD + (1|PAT) + (1|CH), where VOICE (Mother, Stranger) was added as a fixed factor.

**Omnibus tests.** Fixed effects and interactions of each model were assessed with Type II Wald $\chi^2$ tests (car package). As each term was tested once, in a single model including all ROIs, omnibus tests were not corrected for multiple comparisons.

**Post-hoc comparisons.** Post hoc contrasts were computed from estimated marginal means (EMMs) using the emmeans package. They were conducted by ROI when the interaction between the term of interest and ROI was significant (RHY × ROI for Model 1, RHY × FRQ × ROI for Model 2, RHY × FRQ × VOICE × ROI for Model 3), and averaged across ROIs otherwise. As all these interactions were significant, post hoc analyses were conducted by ROI. For model 1, contrasts compared RHY conditions within each ROI. For model 2, they compared RHY conditions at each FRQ level within each ROI. For model 3, two complementary contrasts were derived within each ROI: (a) interaction contrasts between VOICE and RHY within each level of FRQ, and (b) pairwise comparisons of RHY within each combination of FRQ and VOICE. Contrasts were corrected using the multivariate t (mvt) adjustment, equivalent to Tukey's HSD for complete pairwise comparisons, with each family comprising all contrasts within a given set across the eight ROIs (24 contrasts), separately for each frequency band and for each voice in the case of within-voice contrasts. Confidence intervals of contrasts were adjusted accordingly, whereas those of EMMs were not. The exploratory extended model was followed up only because Model 2 produced post hoc contrasts. As its RHY × FRQ × BLK_ORD × ROI interaction was significant, post hoc analyses focused on the estimated slopes (emtrends) of BLK_ORD for each combination of RHY and FRQ within each ROI, tested against zero and compared between RHY conditions, with the same correction. The significance threshold was set at $p < .05$.

**Effect size computation.** Effect sizes were computed for each hypothesis-specific GLMM (Models 1−3) and for the exploratory extended model, following a two-level strategy: model-level and contrast-level effect sizes.

*Model-level effect size.* For each fitted GLMM, marginal $R^2$ ($R^2$m) and conditional $R^2$ ($R^2$c) were computed using the Nakagawa and Schielzeth (*23*) method as implemented in the performance package (r2_nakagawa()). $R^2$m reflects the proportion of variance explained by the fixed effects alone, while $R^2$c reflects the proportion explained by fixed and random effects combined (participant and channel random intercepts). Model log-likelihood was also extracted for each model, and convergence was checked for every model by verifying that the Hessian was positive-definite and that the conditional variance-covariance matrix contained no missing or infinite values, with convergence status logged for all models.

*Contrast-level effect size.* Beyond this model-level quantification, effect sizes for pairwise and interaction contrasts were expressed as unstandardized model-derived estimates on the original scale of the dependent variable (Power, in dB), together with their standard errors and simultaneous 95% confidence intervals (multivariate t adjustment, consistent with the reported *p*-values), all obtained from estimated marginal means (emmeans) computed on the fitted GLMM. Contrasts were computed within each ROI for RHY (Model 1), for RHY within each frequency band (Model 2), and, for Model 3, as RHY × VOICE interaction contrasts within each frequency band and as pairwise RHY contrasts within each combination of frequency band and voice. For the exploratory extended model (Power ~ RHY × FRQ × BLK_ORD × ROI), block-order slopes for all RHY × FRQ conditions were estimated jointly within each ROI, yielding estimated linear trends of Power across blocks (emtrends).

Full model-fit diagnostics ($R^2$m, $R^2$c, log-likelihood, AIC, convergence status) and contrast-level effect sizes are reported in Tables S9–S14.

### 2. Results and Effect sizes

The following tables present statistical results for all eight ROIs (LF, RF, LT, RT, CEN, LP, RP, POS), including electrode clusters not reported in the main text, which focuses on the four frontotemporal clusters (LF, RF, LT, RT). Effect sizes (i.e., model fit statistics [$R^2m$, $R^2c$] and unstandardized model-derived estimates with 95% confidence intervals) are provided in Tables S9–S14.

| Term | Model 1 $\chi^2$(df) [*p*] | Model 2 $\chi^2$(df) [*p*] | Extended model $\chi^2$(df) [*p*] | Model 3 $\chi^2$(df) [*p*] |
|---|---|---|---|---|
| **RHY** | **1054.29** (2) [< .0001] | **1054.33** (2) [< .0001] | **1058.23** (2) [< .0001] | **1057.57** (2) [< .0001] |
| **FRQ** | – | **333.49** (1) [< .0001] | **333.61** (1) [< .0001] | **333.58** (1) [< .0001] |
| **VOICE** | – | – | – | **173.89** (1) [< .0001] |
| **BLK_ORD** | **6.95** (1) [= .0084] | **6.97** (1) [= .0083] | **2.41** (1) [= .1202] | **5.38** (1) [= .0204] |
| **ROI** | **57.62** (7) [< .0001] | **57.62** (7) [< .0001] | **57.59** (7) [< .0001] | **57.57** (7) [< .0001] |
| **RHY × FRQ** | – | **39.28** (2) [< .0001] | **39.71** (2) [< .0001] | **38.94** (2) [< .0001] |
| **RHY × VOICE** | – | – | – | **42.41** (2) [< .0001] |
| **FRQ × VOICE** | – | – | – | **16.56** (1) [< .0001] |
| **RHY × BLK_ORD** | – | – | **215.77** (2) [< .0001] | – |
| **FRQ × BLK_ORD** | – | – | **32.46** (1) [< .0001] | – |
| **RHY × ROI** | **113.60** (14) [< .0001] | **113.66** (14) [< .0001] | **112.61** (14) [< .0001] | **113.78** (14) [< .0001] |
| **FRQ × ROI** | – | **67.42** (7) [< .0001] | **67.37** (7) [< .0001] | **67.48** (7) [< .0001] |
| **VOICE × ROI** | – | – | – | **30.41** (7) [< .0001] |
| **BLK_ORD × ROI** | – | – | **130.07** (7) [< .0001] | – |
| **RHY × FRQ × VOICE** | – | – | – | **85.99** (2) [< .0001] |
| **RHY × FRQ × BLK_ORD** | – | – | **27.76** (2) [< .0001] | – |
| **RHY × FRQ × ROI** | – | **53.46** (14) [< .0001] | **53.73** (14) [< .0001] | **53.69** (14) [< .0001] |
| **RHY × VOICE × ROI** | – | – | – | **71.21** (14) [< .0001] |
| **FRQ × VOICE × ROI** | – | – | – | **16.90** (7) [= .0180] |
| **RHY × BLK_ORD × ROI** | – | – | **55.67** (14) [< .0001] | – |
| **FRQ × BLK_ORD × ROI** | – | – | **18.72** (7) [= .0091] | – |
| **RHY × FRQ × VOICE × ROI** | – | – | – | **36.54** (14) [= .0009] |
| **RHY × FRQ × BLK_ORD × ROI** | – | – | **91.66** (14) [< .0001] | – |

***Note.*** *RHY = rhythmic stimulation (DEL, THE, NR); FRQ = frequency bands (Delta, Theta); VOICE = voice familiarity (Mother, Stranger); ROI = region of interest (LF = Left Frontal; RF = Right Frontal; LT = Left Temporal; RT = Right Temporal; CEN = Central; LP = Left Parietal; RP = Right Parietal; POS = Posterior).*

***Table S4.* Omnibus fixed effects and interactions of the three hypothesis-driven models and of the exploratory extended model.**
Type II Wald $\chi^2$ tests for all terms of each model, reported as $\chi^2$(df) [*p*], without correction for multiple comparisons. All models were fitted to the data of the eight ROIs, with ROI as a fixed factor and random intercepts for participant (PAT) and channel (CH). Model 1: Power ~ RHY × ROI + BLK_ORD. Model 2: Power ~ RHY × FRQ × ROI + BLK_ORD. Model 3: Power ~ RHY × FRQ × VOICE × ROI + BLK_ORD, with BLK_ORD coded as a continuous variable (1–

12). Extended model (exploratory): Power ~ RHY × FRQ × BLK_ORD × ROI, with BLK_ORD realigned within each rhythmic structure (1–4). – = term not included in the model.

| ROI | DEL vs. NR<br>est. [z, *p*] | DEL vs. THE<br>est. [z, *p*] | THE vs. NR<br>est. [z, *p*] |
|---|---|---|---|
| **LF** | **0.227** [13.74, < .0001] | **0.123** [7.53, < .0001] | **0.104** [6.30, < .0001] |
| **RF** | **0.255** [16.25, < .0001] | **0.083** [5.35, < .0001] | **0.171** [10.99, < .0001] |
| **LT** | **0.218** [11.62, < .0001] | *n.s.* [= .4508] | **0.177** [9.47, < .0001] |
| **RT** | **0.268** [14.31, < .0001] | **0.072** [3.85, = .0028] | **0.197** [10.53, < .0001] |
| **CEN** | **0.163** [9.85, < .0001] | **0.070** [4.23, = .0006] | **0.094** [5.67, < .0001] |
| **LP** | **0.097** [4.77, < .0001] | *n.s.* [= .0768] | **0.155** [7.70, < .0001] |
| **RP** | **0.128** [6.30, < .0001] | *n.s.* [= .9999] | **0.144** [7.15, < .0001] |
| **POS** | **0.214** [10.53, < .0001] | *n.s.* [= .0713] | **0.154** [7.63, < .0001] |

***Note.*** *ROI = region of interest (LF = Left Frontal; RF = Right Frontal; LT = Left Temporal; RT = Right Temporal; CEN = Central; LP = Left Parietal; RP = Right Parietal; POS = Posterior); DEL = delta rhythmic stimulation; THE = theta rhythmic stimulation; NR = non-rhythmic stimulation; est. = estimate; n.s. = non-significant.*

***Table S5.*** **Post hoc pairwise contrasts for the RHY factor in all regions of interest.** Contrast estimates reflect marginal mean differences on the response scale; *z*-values and *p*-values are extracted from post hoc tests with multivariate t (mvt) adjustment across the 24 contrasts of the eight ROIs.

| | **Delta** | | | **Theta** | | |
|---|---|---|---|---|---|---|
| **ROI** | **DEL vs. NR**<br>**est.** [*z*, *p*] | **DEL vs. THE**<br>**est.** [*z*, *p*] | **THE vs. NR**<br>**est.** [z, *p*] | **THE vs. DEL**<br>**est.** [*z*, *p*] | **THE vs. NR**<br>**est.** [*z*, *p*] | **DEL vs. NR**<br>**est.** [*z*, *p*] |
| **LF** | **0.138** [5.89, < .0001] | **0.080** [3.43, = .0137] | *n.s.* [= .2397] | **−0.167** [−7.21, < .0001] | **0.149** [6.41, < .0001] | **0.316** [13.54, < .0001] |
| **RF** | **0.215** [9.71, < .0001] | **0.073** [3.33, = .0201] | **0.142** [6.44, < .0001] | **−0.093** [−4.25, = .0005] | **0.201** [9.11, < .0001] | **0.294** [13.28, < .0001] |
| **LT** | **0.119** [4.50, = .0002] | *n.s.* [> .9999] | **0.137** [5.19, < .0001] | **−0.099** [−3.76, = .0039] | **0.217** [8.22, < .0001] | **0.316** [11.92, < .0001] |
| **RT** | **0.246** [9.29, < .0001] | *n.s.* [= .0743] | **0.169** [6.42, < .0001] | *n.s.* [= .2276] | **0.224** [8.49, < .0001] | **0.290** [10.95, < .0001] |
| **CEN** | **0.088** [3.74, = .0042] | *n.s.* [= .9992] | *n.s.* [= .1131] | **−0.117** [−5.01, < .0001] | **0.122** [5.23, < .0001] | **0.239** [10.18, < .0001] |
| **LP** | **0.132** [4.61, < .0001] | *n.s.* [= .8419] | **0.180** [6.32, < .0001] | *n.s.* [= .2812] | **0.130** [4.56, = .0001] | *n.s.* [= .4993] |
| **RP** | **0.120** [4.20, = .0006] | *n.s.* [= .9997] | **0.146** [5.11, < .0001] | *n.s.* [> .9999] | **0.143** [5.01, < .0001] | **0.135** [4.72, < .0001] |
| **POS** | **0.238** [8.30, < .0001] | *n.s.* [= .1053] | **0.158** [5.53, < .0001] | *n.s.* [= .9750] | **0.151** [5.27, < .0001] | **0.189** [6.59, < .0001] |

***Note.*** *Frequency bands (FRQ: Delta, Theta); ROI = region of interest (LF = Left Frontal; RF = Right Frontal; LT = Left Temporal; RT = Right Temporal; CEN = Central; LP = Left Parietal; RP = Right Parietal; POS = Posterior); DEL = delta rhythmic stimulation; THE = theta rhythmic stimulation; NR = non-rhythmic stimulation; est. = estimate; n.s. = non-significant.*

***Table S6.* Post hoc pairwise contrasts for the RHY × FRQ interaction in all regions of interest.**
Contrast estimates reflect marginal mean differences on the response scale; *z*-values and *p*-values extracted from post hoc tests with multivariate t (mvt) adjustment across the 24 contrasts of the eight ROIs, separately for each frequency band. In each band, contrasts are ordered as in the main text (frequency-specificity criteria first), followed by the remaining contrasts.

| ROI | FRQ | Slope DEL est. [*z*, *p*] | Slope NR est. [*z*, *p*] | Slope THE est. [*z*, *p*] | DEL vs. NR est. [*z*, *p*] | DEL vs. THE est. [*z*, *p*] | THE vs. NR est. [*z*, *p*] |
|---|---|---|---|---|---|---|---|
| **LF** | **Delta** | **0.068** [4.36, = .0003] | *n.s.* [= .0652] | *n.s.* [= .0630] | *n.s.* [= .9985] | **0.113** [5.22, < .0001] | **−0.090** [−4.23, = .0005] |
| | **Theta** | *n.s.* [= .8347] | **−0.049** [−3.23, = .0289] | *n.s.* [= .2900] | **0.077** [3.55, = .0091] | *n.s.* [= .0598] | *n.s.* [> .9999] |
| **RF** | **Delta** | **0.111** [7.51, < .0001] | *n.s.* [= .3566] | *n.s.* [> .9999] | **0.077** [3.72, = .0047] | **0.104** [5.09, < .0001] | *n.s.* [= .9739] |
| | **Theta** | **0.074** [5.03, < .0001] | *n.s.* [> .9999] | **0.051** [3.58, = .0083] | **0.067** [3.23, = .0278] | *n.s.* [= .9954] | *n.s.* [= .4939] |
| **LT** | **Delta** | **0.079** [4.48, = .0002] | *n.s.* [= .9715] | *n.s.* [> .9999] | *n.s.* [= .4712] | **0.091** [3.73, = .0045] | *n.s.* [= .9124] |
| | **Theta** | n.s. [> .9999] | *n.s.* [> .9999] | **−0.084** [−4.95, < .0001] | *n.s.* [> .9999] | **0.084** [3.44, = .0132] | **−0.091** [−3.75, = .0042] |
| **RT** | **Delta** | **0.062** [3.53, = .0099] | *n.s.* [= .9997] | *n.s.* [= .0786] | *n.s.* [= .7851] | **0.112** [4.58, = .0001] | *n.s.* [= .1052] |
| | **Theta** | *n.s.* [> .9999] | *n.s.* [= .5047] | **0.053** [3.13, = .0410] | *n.s.* [= .9692] | *n.s.* [= .3503] | **0.091** [3.75, = .0041] |
| **CEN** | **Delta** | **0.078** [4.97, < .0001] | *n.s.* [> .9999] | *n.s.* [= .3716] | **0.071** [3.24, = .0272] | **0.113** [5.21, < .0001] | *n.s.* [= .6402] |
| | **Theta** | **0.089** [5.67, < .0001] | *n.s.* [> .9999] | **−0.076** [−5.07, < .0001] | **0.074** [3.40, = .0156] | **0.165** [7.61, < .0001] | **−0.091** [−4.24, = .0005] |
| **LP** | **Delta** | **0.114** [5.98, < .0001] | *n.s.* [= .9981] | *n.s.* [= .2297] | **0.137** [5.12, < .0001] | **0.161** [6.08, < .0001] | *n.s.* [= .9995] |
| | **Theta** | *n.s.* [= .8791] | *n.s.* [> .9999] | *n.s.* [= .2896] | *n.s.* [= .8970] | *n.s.* [= .0700] | *n.s.* [= .9734] |
| **RP** | **Delta** | *n.s.* [> .9999] | *n.s.* [= .9849] | **−0.097** [−5.28, < .0001] | *n.s.* [= .9779] | **0.107** [4.02, = .0014] | *n.s.* [= .1389] |
| | **Theta** | **−0.100** [−5.23, < .0001] | *n.s.* [= .4607] | **−0.111** [−6.02, < .0001] | *n.s.* [= .4671] | *n.s.* [> .9999] | *n.s.* [= .1694] |
| **POS** | **Delta** | **0.132** [6.88, < .0001] | *n.s.* [= .4181] | *n.s.* [= .9973] | **0.174** [6.52, < .0001] | **0.154** [5.81, < .0001] | *n.s.* [> .9999] |
| | **Theta** | *n.s.* [= .6737] | *n.s.* [= .6914] | **−0.101** [−5.50, < .0001] | *n.s.* [= .1037] | *n.s.* [= .3114] | **−0.138** [−5.28, < .0001] |

***Note.*** *ROI = region of interest (LF = Left Frontal; RF = Right Frontal; LT = Left Temporal; RT = Right Temporal; CEN = Central; LP = Left Parietal; RP = Right Parietal; POS = Posterior); FRQ = frequency bands (Delta, Theta); DEL = delta rhythmic stimulation; THE = theta rhythmic stimulation; NR = non-rhythmic stimulation; est. = estimate; n.s. = non-significant.*

***Table S7.*** **Exploratory analysis: block-order (BLK_ORD) slopes and their pairwise comparisons by rhythmic stimulation (RHY) and frequency band (FRQ) across all regions of interest.**

Slopes and slope differences on the response scale; *z*-values and *p*-values are extracted from post hoc tests with multivariate t (mvt) adjustment across the eight ROIs (24 slopes or contrasts per family, separately for each frequency band).

(1)

| VOICE | **Mother – Stranger** | | | | | |
|---|---|---|---|---|---|---|
| **FRQ** | **Delta** | | | **Theta** | | |
| **ROI** | **DEL vs. NR est.** [z, *p*] | **DEL vs. THE est.** [z, p] | **THE vs. NR est. [z, *p*]** | **DEL vs. NR est.** [z, *p*] | **DEL vs. THE est.** [z, *p*] | **THE vs. NR est.** [z, *p*] |
| **LF** | *n.s.* [> .9999] | *n.s.* [= .6696] | *n.s.* [= .9035] | **−0.291** [−6.22, < .0001] | *n.s.* [= .8193] | **−0.210** [−4.52, = .0001] |
| **RF** | *n.s.* [> .9999] | *n.s.* [> .9999] | *n.s.* [> .9999] | **−0.282** [−6.35, < .0001] | **−0.169** [−3.84, = .0029] | *n.s.* [= .2044] |
| **LT** | *n.s.* [> .9999] | *n.s.* [= .6168] | *n.s.* [= .9884] | **−0.235** [−4.44, = .0002] | **−0.311** [−5.90, < .0001] | *n.s.* [= .9566] |
| **RT** | *n.s.* [= .1232] | **0.170** [3.22, = .0286] | *n.s.* [> .9999] | *n.s.* [= .9995] | *n.s.* [= .0508] | *n.s.* [= .5334] |
| **CEN** | *n.s.* [= .1632] | **0.190** [4.07, = .0011] | *n.s.* [= .9641] | *n.s.* [= .4947] | *n.s.* [> .9999] | *n.s.* [= .5421] |
| **LP** | *n.s.* [> .9999] | *n.s.* [= .9672] | *n.s.* [= .9995] | **−0.176** [−3.07, = .0472] | *n.s.* [= .9868] | *n.s.* [= .7525] |
| **RP** | *n.s.* [= .2035] | *n.s.* [= .9998] | *n.s.* [= .8393] | **−0.216** [−3.76, = .0040] | **−0.209** [−3.67, = .0057] | *n.s.* [> .9999] |
| **POS** | *n.s.* [= .4886] | *n.s.* [= .9946] | *n.s.* [= .9989] | **−0.177** [−3.09, = .0448] | *n.s.* [= .9634] | **−0.257** [−4.50, = .0002] |

(2)

| FRQ | **Delta** | | | | | |
|---|---|---|---|---|---|---|
| **VOICE** | **Mother** | | | **Stranger** | | |
| **ROI** | **DEL vs. NR est.** [z, *p*] | **DEL vs. THE est.** [z, *p*] | **THE vs. NR est.** [z, *p*] | **DEL vs. NR est.** [z, *p*] | **DEL vs. THE est.** [z, *p*] | **THE vs. NR est.** [z, *p*] |
| **LF** | **0.146** [4.53, = .0001] | **0.126** [3.91, = .0022] | *n.s.* [> .9999] | **0.130** [3.84, = .0029] | *n.s.* [= .9969] | *n.s.* [= .1049] |
| **RF** | **0.222** [7.26, < .0001] | *n.s.* [= .0716] | **0.132** [4.29, = .0004] | **0.208** [6.47, < .0001] | *n.s.* [= .7586] | **0.150** [4.76, < .0001] |
| **LT** | **0.139** [3.78, = .0036] | *n.s.* [= .9993] | *n.s.* [= .1099] | *n.s.* [= .1939] | *n.s.* [= .7394] | **0.169** [4.48, = .0002] |
| **RT** | **0.316** [8.63, < .0001] | **0.160** [4.35, = .0003] | **0.156** [4.23, = .0005] | **0.170** [4.43, = .0002] | *n.s.* [> .9999] | **0.180** [4.77, < .0001] |
| **CEN** | **0.148** [4.56, = .0001] | **0.117** [3.62, = .0068] | *n.s.* [= .9996] | *n.s.* [> .9999] | *n.s.* [= .4808] | *n.s.* [= .0885] |
| **LP** | **0.145** [3.67, = .0056] | *n.s.* [> .9999] | **0.152** [3.81, = .0032] | *n.s.* [= .0845] | *n.s.* [= .5297] | **0.206** [5.04, < .0001] |
| **RP** | *n.s.* [= .9810] | *n.s.* [= .9963] | *n.s.* [= .2959] | **0.199** [4.79, < .0001] | *n.s.* [> .9999] | **0.193** [4.74, < .0001] |
| **POS** | **0.298** [7.52, < .0001] | *n.s.* [= .0882] | **0.184** [4.60, = .0001] | **0.174** [4.20, = .0006] | *n.s.* [= .9940] | **0.127** [3.10, = .0435] |

(3)

| FRQ | **Theta** | | | | | |
|---|---|---|---|---|---|---|
| **VOICE** | **Mother** | | | **Stranger** | | |
| **ROI** | **DEL vs. NR est.** [z, *p*] | **DEL vs. THE est.** [z, *p*] | **THE vs. NR est.** [z, *p*] | **DEL vs. NR est.** [z, *p*] | **DEL vs. THE est.** [z, *p*] | **THE vs. NR est.** [z, *p*] |

| | | | | | | |
|---|---|---|---|---|---|---|
| **LF** | **0.179** [5.54, < .0001] | **0.133** [4.12, = .0009] | *n.s.* [= .9644] | **0.469** [13.87, < .0001] | **0.213** [6.41, < .0001] | **0.256** [7.70, < .0001] |
| **RF** | **0.160** [5.24, < .0001] | *n.s.* [> .9999] | **0.147** [4.75, < .0001] | **0.442** [13.78, < .0001] | **0.183** [5.79, < .0001] | **0.260** [8.23, < .0001] |
| **LT** | **0.205** [5.60, < .0001] | *n.s.* [= .9752] | **0.254** [6.88, < .0001] | **0.440** [11.48, < .0001] | **0.262** [6.92, < .0001] | **0.179** [4.74, < .0001] |
| **RT** | **0.314** [8.56, < .0001] | **0.144** [3.94, = .0019] | 0.169 [4.58, = .0001] | **0.264** [6.87, < .0001] | *n.s.* [> .9999] | **0.280** [7.42, < .0001] |
| **CEN** | **0.191** [5.91, < .0001] | **0.118** [3.64, = .0064] | *n.s.* [= .4051] | **0.292** [8.59, < .0001] | **0.121** [3.61, = .0070] | **0.171** [5.12, < .0001] |
| **LP** | *n.s.* [> .9999] | *n.s.* [= .2087] | *n.s.* [= .6357] | **0.154** [3.71, = .0049] | *n.s.* [> .9999] | **0.183** [4.49, = .0002] |
| **RP** | *n.s.* [= .9999] | *n.s.* [= .1306] | **0.141** [3.52, = .0098] | **0.248** [5.98, < .0001] | *n.s.* [= .2592] | **0.148** [3.61, = .0070] |
| **POS** | *n.s.* [= .1672] | *n.s.* [= .5821] | *n.s.* [> .9999] | **0.282** [6.78, < .0001] | *n.s.* [> .9999] | **0.281** [6.88, < .0001] |

***Note.*** *VOICE = voice familiarity (Mother, Stranger); FRQ = frequency bands (Delta, Theta); ROI = region of interest (LF = Left Frontal; RF = Right Frontal; LT = Left Temporal; RT = Right Temporal; CEN = Central; LP = Left Parietal; RP = Right Parietal; POS = Posterior); DEL = delta rhythmic stimulation; THE = theta rhythmic stimulation; NR = non-rhythmic stimulation; est. = estimate; n.s. = non-significant.*

***Table S8.*** **Post hoc contrasts for voice familiarity in all regions of interest: (1) Mother – Stranger interaction contrasts for pairwise rhythm differences in the delta and theta frequency bands, and pairwise rhythm contrasts within each voice in the (2) delta and (3) theta frequency bands.**

Contrast estimates reflect marginal mean differences on the response scale; *z*-values and *p*-values are extracted from post hoc tests with multivariate t (mvt) adjustment across the eight ROIs (24 contrasts per family).

| Model | N observations | $R^2$m | $R^2$c | Log-likelihood | AIC | Convergence |
|---|---|---|---|---|---|---|
| Model 1 | 6,593,544 | 0.00021 | 0.00104 | −21,885,639 | 43,771,335 | Yes |
| Model 2 | 6,593,544 | 0.00028 | 0.00111 | −21,885,392 | 43,770,889 | Yes |
| Extended model (exploratory) | 6,593,544 | 0.00037 | 0.00119 | −21,885,109 | 43,770,415 | Yes |
| Model 3 | 6,593,544 | 0.00035 | 0.00118 | −21,885,155 | 43,770,511 | Yes |

***Table S9.*** **Model fit of the three hypothesis-driven models and of the exploratory extended model: marginal and conditional $R^2$, log-likelihood, AIC, and convergence status (all ROIs, ROI as a factor).**
Model adequacy was assessed via marginal and conditional $R^2$ ($R^2$m, $R^2$c; *23*), which respectively capture the proportion of Power variance attributable to the fixed effects alone vs. to the combined fixed and random-effects structure (including block order, participant and channel random intercepts). Log-likelihood values, AIC and convergence status are reported for each model. Model 1: RHY × ROI. Model 2: RHY × FRQ × ROI. Model 3: RHY × FRQ × VOICE × ROI. Extended model: RHY × FRQ × BLK_ORD × ROI. The low $R^2$ values are expected for single-epoch EEG data, in which trial-to-trial variability is large.

| ROI | Contrast | est. | SE | 95% CI lower | 95% CI upper |
|---|---|---|---|---|---|
| **LF** | DEL vs. NR | 0.227 | 0.017 | 0.176 | 0.277 |
| | DEL vs. THE | 0.123 | 0.016 | 0.073 | 0.173 |
| | THE vs. NR | 0.104 | 0.016 | 0.053 | 0.154 |
| **RF** | DEL vs. NR | 0.255 | 0.016 | 0.207 | 0.303 |
| | DEL vs. THE | 0.083 | 0.016 | 0.036 | 0.131 |
| | THE vs. NR | 0.171 | 0.016 | 0.124 | 0.219 |
| **LT** | DEL vs. NR | 0.218 | 0.019 | 0.160 | 0.275 |
| | DEL vs. THE | 0.041 | 0.019 | −0.016 | 0.098 |
| | THE vs. NR | 0.177 | 0.019 | 0.120 | 0.234 |
| **RT** | DEL vs. NR | 0.268 | 0.019 | 0.211 | 0.326 |
| | DEL vs. THE | 0.072 | 0.019 | 0.015 | 0.128 |
| | THE vs. NR | 0.197 | 0.019 | 0.140 | 0.254 |
| **CEN** | DEL vs. NR | 0.163 | 0.017 | 0.113 | 0.214 |
| | DEL vs. THE | 0.070 | 0.016 | 0.019 | 0.120 |
| | THE vs. NR | 0.094 | 0.017 | 0.043 | 0.144 |
| **LP** | DEL vs. NR | 0.097 | 0.020 | 0.035 | 0.159 |
| | DEL vs. THE | −0.059 | 0.020 | −0.120 | 0.003 |
| | THE vs. NR | 0.155 | 0.020 | 0.094 | 0.217 |
| **RP** | DEL vs. NR | 0.128 | 0.020 | 0.066 | 0.190 |
| | DEL vs. THE | −0.017 | 0.020 | −0.078 | 0.045 |
| | THE vs. NR | 0.144 | 0.020 | 0.083 | 0.206 |
| **POS** | DEL vs. NR | 0.214 | 0.020 | 0.152 | 0.276 |
| | DEL vs. THE | 0.059 | 0.020 | −0.002 | 0.121 |
| | THE vs. NR | 0.154 | 0.020 | 0.093 | 0.216 |

***Note.*** *ROI = region of interest (LF = Left Frontal; RF = Right Frontal; LT = Left Temporal; RT = Right Temporal; CEN = Central; LP = Left Parietal; RP = Right Parietal; POS = Posterior); DEL = delta rhythmic stimulation; THE = theta rhythmic stimulation; NR = non-rhythmic stimulation; est. = unstandardized contrast estimate.*

***Table S10.*** **Post hoc pairwise contrasts for the main effect of the rhythmic stimulation (RHY): estimates, standard errors, and 95% confidence intervals in all regions of interest.**

Pairwise contrast estimates (est.), standard errors (SE), and simultaneous 95% confidence intervals (multivariate t adjustment, consistent with the reported $p$-values), derived via estimated marginal means (EMMs) from the Model 1 generalized linear mixed model predicting Power as a function of rhythmic stimulation (RHY), controlling for block order (BLK_ORD), with random intercepts for participant and channel. Contrasts compare the rhythmic structure of the stimuli (DEL, THE, NR) and reflect raw differences in estimated marginal means on the response scale. $z$- and mvt-adjusted $p$-values are reported in Table S5.

| ROI | FRQ | Contrast | est. | SE | 95% CI lower | 95% CI upper |
|---|---|---|---|---|---|---|
| **LF** | Delta | DEL vs. NR | 0.138 | 0.023 | 0.066 | 0.209 |
| | | DEL vs. THE | 0.080 | 0.023 | 0.009 | 0.150 |
| | | THE vs. NR | 0.058 | 0.023 | −0.013 | 0.129 |
| | Theta | THE vs. DEL | −0.167 | 0.023 | −0.238 | −0.096 |
| | | THE vs. NR | 0.149 | 0.023 | 0.078 | 0.220 |
| | | DEL vs. NR | 0.316 | 0.023 | 0.245 | 0.387 |
| **RF** | Delta | DEL vs. NR | 0.215 | 0.022 | 0.147 | 0.283 |
| | | DEL vs. THE | 0.073 | 0.022 | 0.006 | 0.140 |
| | | THE vs. NR | 0.142 | 0.022 | 0.075 | 0.209 |
| | Theta | THE vs. DEL | −0.093 | 0.022 | −0.161 | −0.026 |
| | | THE vs. NR | 0.201 | 0.022 | 0.134 | 0.268 |
| | | DEL vs. NR | 0.294 | 0.022 | 0.227 | 0.362 |
| **LT** | Delta | DEL vs. NR | 0.119 | 0.026 | 0.038 | 0.200 |
| | | DEL vs. THE | −0.018 | 0.026 | −0.098 | 0.063 |
| | | THE vs. NR | 0.137 | 0.026 | 0.056 | 0.218 |
| | Theta | THE vs. DEL | −0.099 | 0.026 | −0.179 | −0.019 |
| | | THE vs. NR | 0.217 | 0.026 | 0.136 | 0.297 |
| | | DEL vs. NR | 0.316 | 0.026 | 0.235 | 0.397 |
| **RT** | Delta | DEL vs. NR | 0.246 | 0.027 | 0.165 | 0.327 |
| | | DEL vs. THE | 0.077 | 0.026 | −0.003 | 0.157 |
| | | THE vs. NR | 0.169 | 0.026 | 0.089 | 0.250 |
| | Theta | THE vs. DEL | −0.066 | 0.026 | −0.147 | 0.014 |
| | | THE vs. NR | 0.224 | 0.026 | 0.144 | 0.305 |
| | | DEL vs. NR | 0.290 | 0.027 | 0.209 | 0.371 |
| **CEN** | Delta | DEL vs. NR | 0.088 | 0.023 | 0.016 | 0.159 |
| | | DEL vs. THE | 0.023 | 0.023 | −0.048 | 0.094 |
| | | THE vs. NR | 0.065 | 0.023 | −0.006 | 0.136 |
| | Theta | THE vs. DEL | −0.117 | 0.023 | −0.188 | −0.045 |
| | | THE vs. NR | 0.122 | 0.023 | 0.051 | 0.193 |
| | | DEL vs. NR | 0.239 | 0.023 | 0.167 | 0.310 |
| **LP** | Delta | DEL vs. NR | 0.132 | 0.029 | 0.045 | 0.220 |
| | | DEL vs. THE | −0.048 | 0.028 | −0.135 | 0.039 |
| | | THE vs. NR | 0.180 | 0.029 | 0.093 | 0.268 |
| | Theta | THE vs. DEL | 0.069 | 0.028 | −0.018 | 0.156 |
| | | THE vs. NR | 0.130 | 0.029 | 0.043 | 0.217 |
| | | DEL vs. NR | 0.061 | 0.029 | −0.026 | 0.149 |
| **RP** | Delta | DEL vs. NR | 0.120 | 0.029 | 0.033 | 0.208 |
| | | DEL vs. THE | −0.026 | 0.028 | −0.113 | 0.061 |
| | | THE vs. NR | 0.146 | 0.029 | 0.059 | 0.233 |
| | Theta | THE vs. DEL | 0.008 | 0.028 | −0.079 | 0.095 |
| | | THE vs. NR | 0.143 | 0.029 | 0.056 | 0.230 |
| | | DEL vs. NR | 0.135 | 0.029 | 0.048 | 0.223 |
| **POS** | Delta | DEL vs. NR | 0.238 | 0.029 | 0.150 | 0.326 |
| | | DEL vs. THE | 0.080 | 0.028 | −0.007 | 0.167 |
| | | THE vs. NR | 0.158 | 0.029 | 0.071 | 0.245 |
| | Theta | THE vs. DEL | −0.038 | 0.028 | −0.125 | 0.049 |

| | | | | | | |
|---|---|---|---|---|---|---|
| | | THE vs. NR | 0.151 | 0.029 | 0.063 | 0.238 |
| | | DEL vs. NR | 0.189 | 0.029 | 0.101 | 0.277 |

***Note.*** *ROI = region of interest (LF = Left Frontal; RF = Right Frontal; LT = Left Temporal; RT = Right Temporal; CEN = Central; LP = Left Parietal; RP = Right Parietal; POS = Posterior); FRQ = frequency bands (Delta, Theta); DEL = delta rhythmic stimulation; THE = theta rhythmic stimulation; NR = non-rhythmic stimulation; est. = unstandardized contrast estimate.*

***Table S11.*** **Post hoc pairwise contrasts for the RHY × FRQ interaction: estimates, standard errors, and 95% confidence intervals by frequency band in all regions of interest.**

Pairwise contrast estimates (est.), standard errors (SE), and simultaneous 95% confidence intervals (multivariate t adjustment, consistent with the reported *p*-values), derived via estimated marginal means (EMMs) from the Model 2 generalized linear mixed model predicting Power as a function of rhythmic stimulation (RHY), frequency band (FRQ), and their interaction (RHY × FRQ), controlling for block order (BLK_ORD), with random intercepts for participant and channel. Contrasts compare the rhythmic structure of the stimuli (DEL, THE, NR) within each frequency band (Delta, Theta) and reflect raw differences in estimated marginal means on the response scale. *z*- and mvt-adjusted *p*-values are reported in Table S6.

| ROI | FRQ | RHY | Slope est. | SE | 95% CI lower | 95% CI upper |
|---|---|---|---|---|---|---|
| **LF** | Delta | DEL | 0.068 | 0.016 | 0.020 | 0.116 |
| | | NR | 0.045 | 0.015 | −0.001 | 0.092 |
| | | THE | −0.045 | 0.015 | −0.091 | 0.001 |
| | Theta | DEL | 0.028 | 0.016 | −0.020 | 0.076 |
| | | NR | −0.049 | 0.015 | −0.096 | −0.002 |
| | | THE | −0.037 | 0.015 | −0.083 | 0.009 |
| **RF** | Delta | DEL | 0.111 | 0.015 | 0.066 | 0.156 |
| | | NR | 0.034 | 0.014 | −0.010 | 0.078 |
| | | THE | 0.007 | 0.014 | −0.037 | 0.050 |
| | Theta | DEL | 0.074 | 0.015 | 0.029 | 0.120 |
| | | NR | 0.007 | 0.014 | −0.037 | 0.052 |
| | | THE | 0.051 | 0.014 | 0.007 | 0.095 |
| **LT** | Delta | DEL | 0.079 | 0.018 | 0.025 | 0.133 |
| | | NR | 0.026 | 0.017 | −0.027 | 0.079 |
| | | THE | −0.012 | 0.017 | −0.064 | 0.040 |
| | Theta | DEL | 0.000 | 0.018 | −0.054 | 0.055 |
| | | NR | 0.007 | 0.017 | −0.046 | 0.060 |
| | | THE | −0.084 | 0.017 | −0.136 | −0.032 |
| **RT** | Delta | DEL | 0.062 | 0.018 | 0.008 | 0.117 |
| | | NR | 0.018 | 0.017 | −0.035 | 0.071 |
| | | THE | −0.050 | 0.017 | −0.102 | 0.002 |
| | Theta | DEL | −0.004 | 0.018 | −0.058 | 0.050 |
| | | NR | −0.038 | 0.017 | −0.091 | 0.015 |
| | | THE | 0.053 | 0.017 | 0.001 | 0.105 |
| **CEN** | Delta | DEL | 0.078 | 0.016 | 0.030 | 0.126 |
| | | NR | 0.007 | 0.015 | −0.040 | 0.054 |
| | | THE | −0.035 | 0.015 | −0.081 | 0.011 |
| | Theta | DEL | 0.089 | 0.016 | 0.041 | 0.137 |
| | | NR | 0.015 | 0.015 | −0.032 | 0.061 |
| | | THE | −0.076 | 0.015 | −0.123 | −0.030 |
| **LP** | Delta | DEL | 0.114 | 0.019 | 0.056 | 0.173 |
| | | NR | −0.022 | 0.019 | −0.080 | 0.035 |
| | | THE | −0.047 | 0.018 | −0.103 | 0.010 |
| | Theta | DEL | 0.033 | 0.019 | −0.026 | 0.092 |
| | | NR | −0.010 | 0.019 | −0.067 | 0.048 |
| | | THE | −0.045 | 0.018 | −0.102 | 0.011 |
| **RP** | Delta | DEL | 0.009 | 0.019 | −0.049 | 0.068 |
| | | NR | −0.026 | 0.019 | −0.084 | 0.031 |
| | | THE | −0.097 | 0.018 | −0.154 | −0.041 |
| | Theta | DEL | −0.100 | 0.019 | −0.158 | −0.041 |
| | | NR | −0.042 | 0.019 | −0.099 | 0.016 |
| | | THE | −0.111 | 0.018 | −0.167 | −0.054 |
| **POS** | Delta | DEL | 0.132 | 0.019 | 0.073 | 0.190 |
| | | NR | −0.043 | 0.019 | −0.100 | 0.015 |
| | | THE | −0.023 | 0.018 | −0.079 | 0.034 |
| | Theta | DEL | −0.038 | 0.019 | −0.097 | 0.020 |

| | | | | | | |
|---|---|---|---|---|---|---|
| | | NR | 0.037 | 0.019 | −0.020 | 0.094 |
| | | THE | −0.101 | 0.018 | −0.158 | −0.045 |

***Note.*** *ROI = region of interest (LF = Left Frontal; RF = Right Frontal; LT = Left Temporal; RT = Right Temporal; CEN = Central; LP = Left Parietal; RP = Right Parietal; POS = Posterior); DEL = delta rhythmic stimulation; THE = theta rhythmic stimulation; NR = non-rhythmic stimulation; FRQ = frequency bands (Delta, Theta); est. = unstandardized slope estimate.*

***Table S12.*** **Estimated marginal slopes of block order (BLK_ORD), standard error, and 95% confidence intervals from the mixed-effects model by condition in all regions of interest.**

Slope estimates (est.), standard errors (SE), and simultaneous 95% confidence intervals (multivariate t adjustment, consistent with the reported *p*-values), derived via emtrends from the extended Model 2 generalized linear mixed model with random intercepts for participant and channel, predicting Power as a function of block order (BLK_ORD) with a full RHY × FRQ × BLK_ORD interaction. Marginal slopes were extracted separately for each RHY × FRQ condition. Slopes reflect the estimated linear trend in Power across experimental blocks and are tested against the null hypothesis (slope = 0). *z*- and mvt-adjusted *p*-values are reported in Table S7.

| ROI | FRQ | Contrast (Mother − Stranger) | est. | SE | 95% CI lower | 95% CI upper |
|---|---|---|---|---|---|---|
| **LF** | Delta | DEL vs. NR | 0.016 | 0.047 | −0.127 | 0.159 |
| | | DEL vs. THE | 0.090 | 0.046 | −0.052 | 0.231 |
| | | THE vs. NR | −0.074 | 0.046 | −0.215 | 0.068 |
| | Theta | DEL vs. NR | −0.291 | 0.047 | −0.433 | −0.148 |
| | | DEL vs. THE | −0.080 | 0.046 | −0.222 | 0.061 |
| | | THE vs. NR | −0.210 | 0.046 | −0.352 | −0.068 |
| **RF** | Delta | DEL vs. NR | 0.015 | 0.044 | −0.121 | 0.150 |
| | | DEL vs. THE | 0.032 | 0.044 | −0.102 | 0.167 |
| | | THE vs. NR | −0.018 | 0.044 | −0.153 | 0.117 |
| | Theta | DEL vs. NR | −0.282 | 0.044 | −0.417 | −0.146 |
| | | DEL vs. THE | −0.169 | 0.044 | −0.303 | −0.034 |
| | | THE vs. NR | −0.113 | 0.044 | −0.248 | 0.022 |
| **LT** | Delta | DEL vs. NR | 0.039 | 0.053 | −0.122 | 0.201 |
| | | DEL vs. THE | 0.105 | 0.053 | −0.056 | 0.266 |
| | | THE vs. NR | −0.066 | 0.053 | −0.227 | 0.096 |
| | Theta | DEL vs. NR | −0.235 | 0.053 | −0.397 | −0.073 |
| | | DEL vs. THE | −0.311 | 0.053 | −0.472 | −0.150 |
| | | THE vs. NR | 0.076 | 0.053 | −0.086 | 0.237 |
| **RT** | Delta | DEL vs. NR | 0.146 | 0.053 | −0.016 | 0.308 |
| | | DEL vs. THE | 0.170 | 0.053 | 0.009 | 0.331 |
| | | THE vs. NR | −0.024 | 0.053 | −0.185 | 0.138 |
| | Theta | DEL vs. NR | 0.050 | 0.053 | −0.112 | 0.212 |
| | | DEL vs. THE | 0.161 | 0.053 | −0.000 | 0.321 |
| | | THE vs. NR | −0.111 | 0.053 | −0.272 | 0.051 |
| **CEN** | Delta | DEL vs. NR | 0.124 | 0.047 | −0.019 | 0.268 |
| | | DEL vs. THE | 0.190 | 0.047 | 0.047 | 0.332 |
| | | THE vs. NR | −0.065 | 0.047 | −0.208 | 0.077 |
| | Theta | DEL vs. NR | −0.100 | 0.047 | −0.244 | 0.043 |
| | | DEL vs. THE | −0.003 | 0.047 | −0.145 | 0.139 |
| | | THE vs. NR | −0.097 | 0.047 | −0.240 | 0.045 |
| **LP** | Delta | DEL vs. NR | 0.026 | 0.057 | −0.150 | 0.201 |
| | | DEL vs. THE | 0.079 | 0.057 | −0.095 | 0.253 |
| | | THE vs. NR | −0.053 | 0.057 | −0.228 | 0.121 |
| | Theta | DEL vs. NR | −0.176 | 0.057 | −0.351 | −0.001 |
| | | DEL vs. THE | −0.072 | 0.057 | −0.246 | 0.102 |
| | | THE vs. NR | −0.104 | 0.057 | −0.279 | 0.070 |
| **RP** | Delta | DEL vs. NR | −0.147 | 0.057 | −0.322 | 0.028 |
| | | DEL vs. THE | −0.050 | 0.057 | −0.224 | 0.124 |
| | | THE vs. NR | −0.097 | 0.057 | −0.272 | 0.077 |
| | Theta | DEL vs. NR | −0.216 | 0.057 | −0.391 | −0.040 |
| | | DEL vs. THE | −0.209 | 0.057 | −0.383 | −0.035 |
| | | THE vs. NR | −0.007 | 0.057 | −0.181 | 0.168 |
| **POS** | Delta | DEL vs. NR | 0.123 | 0.057 | −0.052 | 0.299 |
| | | DEL vs. THE | 0.066 | 0.057 | −0.108 | 0.240 |
| | | THE vs. NR | 0.057 | 0.057 | −0.117 | 0.232 |
| | Theta | DEL vs. NR | −0.177 | 0.057 | −0.353 | −0.002 |

| | | | | | | |
|---|---|---|---|---|---|---|
| | | DEL vs. THE | 0.080 | 0.057 | −0.094 | 0.254 |
| | | THE vs. NR | −0.257 | 0.057 | −0.432 | −0.083 |

***Note.*** *ROI = region of interest (LF = Left Frontal; RF = Right Frontal; LT = Left Temporal; RT = Right Temporal; CEN = Central; LP = Left Parietal; RP = Right Parietal; POS = Posterior); FRQ = frequency bands (Delta, Theta); DEL = delta rhythmic stimulation; THE = theta rhythmic stimulation; NR = non-rhythmic stimulation; est. = unstandardized interaction contrast estimate.*

***Table S13.*** **Between-voice interaction contrasts (Mother − Stranger) for pairwise rhythm differences: estimates, standard errors, and 95% confidence intervals by frequency band in all regions of interest..**

Pairwise contrast estimates (est.), standard errors (SE), and simultaneous 95% confidence intervals (multivariate t adjustment, consistent with the reported *p*-values), derived via estimated marginal means (EMMs) from the Model 3 generalized linear mixed model predicting Power as a function of rhythmic stimulation (RHY), frequency band (FRQ), voice familiarity (VOICE), and their interactions (RHY × FRQ × VOICE), controlling for block order (BLK_ORD), with random intercepts for participant and channel. Contrasts reflect the between-voice difference (Mother−Stranger) in stimulus rhythmic-structure contrasts (DEL, THE, NR) within each frequency band (Delta, Theta) and represent raw differences in estimated marginal means on the response scale. *z*- and mvt-adjusted *p*-values are reported in Table S8, panel 1.

| ROI | VOICE | FRQ | Contrast | est. | SE | 95% CI lower | 95% CI upper |
|---|---|---|---|---|---|---|---|
| **LF** | Mother | Delta | DEL vs. NR | 0.146 | 0.032 | 0.048 | 0.245 |
| | | | DEL vs. THE | 0.126 | 0.032 | 0.028 | 0.225 |
| | | | THE vs. NR | 0.020 | 0.033 | −0.080 | 0.119 |
| | | Theta | DEL vs. NR | 0.179 | 0.032 | 0.080 | 0.277 |
| | | | DEL vs. THE | 0.133 | 0.032 | 0.035 | 0.232 |
| | | | THE vs. NR | 0.046 | 0.033 | −0.054 | 0.145 |
| | Stranger | Delta | DEL vs. NR | 0.130 | 0.034 | 0.027 | 0.233 |
| | | | DEL vs. THE | 0.037 | 0.033 | −0.065 | 0.138 |
| | | | THE vs. NR | 0.093 | 0.033 | −0.008 | 0.195 |
| | | Theta | DEL vs. NR | 0.469 | 0.034 | 0.366 | 0.572 |
| | | | DEL vs. THE | 0.213 | 0.033 | 0.112 | 0.315 |
| | | | THE vs. NR | 0.256 | 0.033 | 0.154 | 0.357 |
| **RF** | Mother | Delta | DEL vs. NR | 0.222 | 0.031 | 0.129 | 0.316 |
| | | | DEL vs. THE | 0.090 | 0.031 | −0.004 | 0.184 |
| | | | THE vs. NR | 0.132 | 0.031 | 0.038 | 0.227 |
| | | Theta | DEL vs. NR | 0.160 | 0.031 | 0.067 | 0.254 |
| | | | DEL vs. THE | 0.014 | 0.031 | −0.080 | 0.107 |
| | | | THE vs. NR | 0.147 | 0.031 | 0.052 | 0.241 |
| | Stranger | Delta | DEL vs. NR | 0.208 | 0.032 | 0.110 | 0.306 |
| | | | DEL vs. THE | 0.058 | 0.032 | −0.039 | 0.154 |
| | | | THE vs. NR | 0.150 | 0.032 | 0.054 | 0.246 |
| | | Theta | DEL vs. NR | 0.442 | 0.032 | 0.344 | 0.540 |
| | | | DEL vs. THE | 0.183 | 0.032 | 0.086 | 0.279 |
| | | | THE vs. NR | 0.260 | 0.032 | 0.163 | 0.356 |
| **LT** | Mother | Delta | DEL vs. NR | 0.139 | 0.037 | 0.027 | 0.250 |
| | | | DEL vs. THE | 0.035 | 0.037 | −0.077 | 0.147 |
| | | | THE vs. NR | 0.103 | 0.037 | −0.010 | 0.216 |
| | | Theta | DEL vs. NR | 0.205 | 0.037 | 0.093 | 0.317 |
| | | | DEL vs. THE | −0.049 | 0.037 | −0.161 | 0.063 |
| | | | THE vs. NR | 0.254 | 0.037 | 0.142 | 0.367 |
| | Stranger | Delta | DEL vs. NR | 0.099 | 0.038 | −0.018 | 0.216 |
| | | | DEL vs. THE | −0.070 | 0.038 | −0.185 | 0.046 |
| | | | THE vs. NR | 0.169 | 0.038 | 0.054 | 0.284 |
| | | Theta | DEL vs. NR | 0.440 | 0.038 | 0.323 | 0.558 |
| | | | DEL vs. THE | 0.262 | 0.038 | 0.146 | 0.377 |
| | | | THE vs. NR | 0.179 | 0.038 | 0.064 | 0.294 |
| **RT** | Mother | Delta | DEL vs. NR | 0.316 | 0.037 | 0.204 | 0.428 |
| | | | DEL vs. THE | 0.160 | 0.037 | 0.048 | 0.272 |
| | | | THE vs. NR | 0.156 | 0.037 | 0.044 | 0.269 |
| | | Theta | DEL vs. NR | 0.314 | 0.037 | 0.202 | 0.426 |
| | | | DEL vs. THE | 0.144 | 0.037 | 0.032 | 0.256 |
| | | | THE vs. NR | 0.169 | 0.037 | 0.057 | 0.282 |
| | Stranger | Delta | DEL vs. NR | 0.170 | 0.038 | 0.053 | 0.287 |
| | | | DEL vs. THE | −0.010 | 0.038 | −0.126 | 0.105 |
| | | | THE vs. NR | 0.180 | 0.038 | 0.065 | 0.295 |
| | | Theta | DEL vs. NR | 0.264 | 0.038 | 0.146 | 0.381 |

| | | | | | | | |
|---|---|---|---|---|---|---|---|
| | | | DEL vs. THE | −0.016 | 0.038 | −0.132 | 0.099 |
| | | | THE vs. NR | 0.280 | 0.038 | 0.165 | 0.395 |
| **CEN** | Mother | Delta | DEL vs. NR | 0.148 | 0.032 | 0.049 | 0.247 |
| | | | DEL vs. THE | 0.117 | 0.032 | 0.018 | 0.216 |
| | | | THE vs. NR | 0.030 | 0.033 | −0.070 | 0.130 |
| | | Theta | DEL vs. NR | 0.191 | 0.032 | 0.093 | 0.290 |
| | | | DEL vs. THE | 0.118 | 0.032 | 0.019 | 0.217 |
| | | | THE vs. NR | 0.074 | 0.033 | −0.026 | 0.173 |
| | Stranger | Delta | DEL vs. NR | 0.023 | 0.034 | −0.080 | 0.127 |
| | | | DEL vs. THE | −0.072 | 0.033 | −0.174 | 0.030 |
| | | | THE vs. NR | 0.096 | 0.033 | −0.006 | 0.198 |
| | | Theta | DEL vs. NR | 0.292 | 0.034 | 0.188 | 0.396 |
| | | | DEL vs. THE | 0.121 | 0.033 | 0.019 | 0.223 |
| | | | THE vs. NR | 0.171 | 0.033 | 0.069 | 0.273 |
| **LP** | Mother | Delta | DEL vs. NR | 0.145 | 0.040 | 0.024 | 0.266 |
| | | | DEL vs. THE | −0.007 | 0.040 | −0.128 | 0.114 |
| | | | THE vs. NR | 0.152 | 0.040 | 0.030 | 0.274 |
| | | Theta | DEL vs. NR | −0.022 | 0.040 | −0.143 | 0.099 |
| | | | DEL vs. THE | −0.101 | 0.040 | −0.222 | 0.020 |
| | | | THE vs. NR | 0.079 | 0.040 | −0.043 | 0.201 |
| | Stranger | Delta | DEL vs. NR | 0.120 | 0.042 | −0.007 | 0.247 |
| | | | DEL vs. THE | −0.086 | 0.041 | −0.211 | 0.039 |
| | | | THE vs. NR | 0.206 | 0.041 | 0.081 | 0.330 |
| | | Theta | DEL vs. NR | 0.154 | 0.042 | 0.027 | 0.281 |
| | | | DEL vs. THE | −0.029 | 0.041 | −0.154 | 0.096 |
| | | | THE vs. NR | 0.183 | 0.041 | 0.059 | 0.308 |
| **RP** | Mother | Delta | DEL vs. NR | 0.052 | 0.040 | −0.069 | 0.173 |
| | | | DEL vs. THE | −0.044 | 0.040 | −0.165 | 0.077 |
| | | | THE vs. NR | 0.096 | 0.040 | −0.026 | 0.218 |
| | | Theta | DEL vs. NR | 0.033 | 0.040 | −0.088 | 0.154 |
| | | | DEL vs. THE | −0.108 | 0.040 | −0.229 | 0.013 |
| | | | THE vs. NR | 0.141 | 0.040 | 0.019 | 0.263 |
| | Stranger | Delta | DEL vs. NR | 0.199 | 0.042 | 0.072 | 0.326 |
| | | | DEL vs. THE | 0.005 | 0.041 | −0.120 | 0.130 |
| | | | THE vs. NR | 0.193 | 0.041 | 0.069 | 0.318 |
| | | Theta | DEL vs. NR | 0.248 | 0.042 | 0.122 | 0.375 |
| | | | DEL vs. THE | 0.101 | 0.041 | −0.024 | 0.226 |
| | | | THE vs. NR | 0.148 | 0.041 | 0.023 | 0.272 |
| **POS** | Mother | Delta | DEL vs. NR | 0.298 | 0.040 | 0.177 | 0.419 |
| | | | DEL vs. THE | 0.114 | 0.040 | −0.007 | 0.235 |
| | | | THE vs. NR | 0.184 | 0.040 | 0.062 | 0.306 |
| | | Theta | DEL vs. NR | 0.105 | 0.040 | −0.016 | 0.226 |
| | | | DEL vs. THE | 0.081 | 0.040 | −0.040 | 0.202 |
| | | | THE vs. NR | 0.024 | 0.040 | −0.098 | 0.146 |
| | Stranger | Delta | DEL vs. NR | 0.174 | 0.042 | 0.048 | 0.301 |
| | | | DEL vs. THE | 0.048 | 0.041 | −0.077 | 0.173 |
| | | | THE vs. NR | 0.127 | 0.041 | 0.002 | 0.251 |

| | | | | | | | |
|---|---|---|---|---|---|---|---|
| | | Theta | DEL vs. NR | 0.282 | 0.042 | 0.155 | 0.409 |
| | | | DEL vs. THE | 0.001 | 0.041 | −0.124 | 0.126 |
| | | | THE vs. NR | 0.281 | 0.041 | 0.156 | 0.406 |

***Note.*** *ROI = region of interest (LF = Left Frontal; RF = Right Frontal; LT = Left Temporal; RT = Right Temporal; CEN = Central; LP = Left Parietal; RP = Right Parietal; POS = Posterior); VOICE = voice familiarity (Mother, Stranger); FRQ = frequency bands (Delta, Theta); DEL = delta rhythmic stimulation; THE = theta rhythmic stimulation; NR = non-rhythmic stimulation; est. = unstandardized contrast estimate.*

***Table S14.*** **Post hoc pairwise contrasts for the rhythmic (RHY) effect within each voice condition: estimates, standard errors, and 95% confidence intervals by frequency band in all regions of interest.**

Pairwise contrast estimates (est.), standard errors (SE), and simultaneous 95% confidence intervals (multivariate t adjustment, consistent with the reported *p*-values), derived via estimated marginal means (EMMs) from the Model 3 generalized linear mixed model predicting Power as a function of stimulus rhythmic stimulation (RHY), frequency band (FRQ), voice familiarity (VOICE), and their interactions (RHY × FRQ × VOICE), controlling for block order (BLK_ORD), with random intercepts for participant and channel. Contrasts compare the rhythmic stimulation (DEL, THE, NR) within each combination of voice condition (Mother, Stranger) and frequency band (Delta, Theta), and reflect raw differences in estimated marginal means on the response scale. *z*- and mvt-adjusted *p*-values are reported in Table S8, panels 2 and 3.

### References

1. L. Murray, A. D. Carothers, The validation of the Edinburgh Post-natal Depression Scale on a community sample. *The British Journal of Psychiatry* **157**, 288-290 (1990).
2. M. A. Gartstein, M. K. Rothbart, Studying infant temperament via the revised infant behavior questionnaire. *Infant behavior and development* **26**, 64-86 (2003).
3. R. R. Abidin, *Parenting Stress Index: Professional Manual* (Psychological Assessment Resources, Odessa, FL, ed. 3, 1995).
4. R. Largo, D. Pfister, L. Molinari, S. Kundu, A. Lipp, G. Due, Significance of prenatal, perinatal and postnatal factors in the development of AGA preterm infants at five to seven years. *Developmental Medicine & Child Neurology* **31**, 440-456 (1989).
5. N. N. Soe, D. J. Wen, J. S. Poh, Y. Li, B. F. Broekman, H. Chen, Y. S. Chong, K. Kwek, S.-M. Saw, P. D. Gluckman, Pre-and post-natal maternal depressive symptoms in relation with infant frontal function, connectivity, and behaviors. *PloS one* **11**, e0152991 (2016).
6. P. Tomalski, D. G. Moore, H. Ribeiro, E. L. Axelsson, E. Murphy, A. Karmiloff-Smith, M. H. Johnson, E. Kushnerenko, Socioeconomic status and functional brain development–associations in early infancy. *Developmental science* **16**, 676-687 (2013).
7. M. M. Crespo-Llado, R. Vanderwert, E. Roberti, E. Geangu, Eight-month-old infants' behavioral responses to peers' emotions as related to the asymmetric frontal cortex activity. *Scientific reports* **8**, 17152 (2018).
8. R Core Team. (R Foundation for Statistical Computing, 2025).
9. M. E. Brooks, K. Kristensen, K. J. van Benthem, A. Magnusson, C. W. Berg, A. Nielsen, H. J. Skaug, M. Maechler, B. M. Bolker, glmmTMB balances speed and flexibility among packages for zero-inflated generalized linear mixed modeling. *The R Journal* **9**, 378-400 (2017).
10. R. V. Lenth, J. Piaskowski. (CRAN, 2025).
11. A. Genz, F. Bretz, *Computation of Multivariate Normal and t Probabilities* (Springer, Berlin, Heidelberg, 2009), vol. 195.
12. J. Fox, S. Weisberg, *An R Companion to Applied Regression* (Sage, Thousand Oaks, CA, ed. 3, 2019).
13. D. Lüdecke, M. S. Ben-Shachar, I. Patil, P. Waggoner, D. Makowski, performance: An R package for assessment, comparison and testing of statistical models. *Journal of Open Source Software* **6**, 3139 (2021).
14. F. Hartig, in *CRAN*. (2024).
15. H. Wickham, *ggplot2: Elegant Graphics for Data Analysis* (Springer, New York, ed. 2, 2016).
16. G. Z. Tau, B. S. Peterson, Normal development of brain circuits. *Neuropsychopharmacology* **35**, 147-168 (2010).
17. P. J. Uhlhaas, F. Roux, E. Rodriguez, A. Rotarska-Jagiela, W. Singer, Neural synchrony and the development of cortical networks. *Trends in cognitive sciences* **14**, 72-80 (2010).
18. M. C. Ortiz-Barajas, R. Guevara, J. Gervain, Neural oscillations and speech processing at birth. *iScience* **26**, (2023).
19. A. Attaheri, D. Panayiotou, A. Phillips, A. Ni Choisdealbha, G. M. Di Liberto, S. Rocha, P. Brusini, N. Mead, S. Flanagan, H. Olawole-Scott, U. Goswami, Cortical

Tracking of Sung Speech in Adults vs Infants: A Developmental Analysis. *Front Neurosci* **16**, 842447 (2022).
20. A. Attaheri, A. Ni Choisdealbha, G. M. Di Liberto, S. Rocha, P. Brusini, N. Mead, H. Olawole-Scott, P. Boutris, S. Gibbon, I. Williams, C. Grey, S. Flanagan, U. Goswami, Delta- and theta-band cortical tracking and phase-amplitude coupling to sung speech by infants. *Neuroimage* **247**, 118698 (2022).
21. P. J. Marshall, Y. Bar-Haim, N. A. Fox, Development of the EEG from 5 months to 4 years of age. *Clinical neurophysiology* **113**, 1199-1208 (2002).
22. L. K. Cirelli, C. Spinelli, S. Nozaradan, L. J. Trainor, Measuring neural entrainment to beat and meter in infants: effects of music background. *Frontiers in neuroscience* **10**, 229 (2016).
23. S. Nakagawa, H. Schielzeth, A general and simple method for obtaining R2 from generalized linear mixed-effects models. *Methods in ecology and evolution* **4**, 133-142 (2013).
